# Probing the Structural Dynamics of the Unbound MAX Protein: Insights from Well-Tempered Metadynamics

**DOI:** 10.1101/2025.09.01.673476

**Authors:** Huixia Lu, Jordi Marti, Jordi Faraudo

## Abstract

One of most relevant challenges in tumorigenesis is the association of MYC and MAX proteins, whose related cancers remain undrugged. In particular, the disordered regions shown by those oncogenes make their structural characterization and the development of new drugs a truly hard task. To address these challenges, we employed the enhanced-sampling well-tempered metadynamics method to systematically explore the conformational space of the unbound MAX protein. Our results revealed, for the first time, a well-defined and thermodynamically favorable conformation of monomeric MAX. This is a remarkable finding, as it demonstrates that regions of MAX previously considered persistently disordered are capable of adopting stable, folded structures under specific conditions. Moreover, our findings also suggest that the meta-stable structural motifs observed in this work may harbor druggable sites, particularly relevant for strategies aiming to target MAX directly or to disrupt its interaction with MYC, thereby modulating oncogenic signaling pathways. The present study establishes a new structural framework for understanding the dynamics of MAX and provides a foundation for future structure-based drug design targeting the MYC/MAX axis. Finally, our work offers a strategic blueprint for investigating similarly challenging drug targets.

**TOC Graphic:** 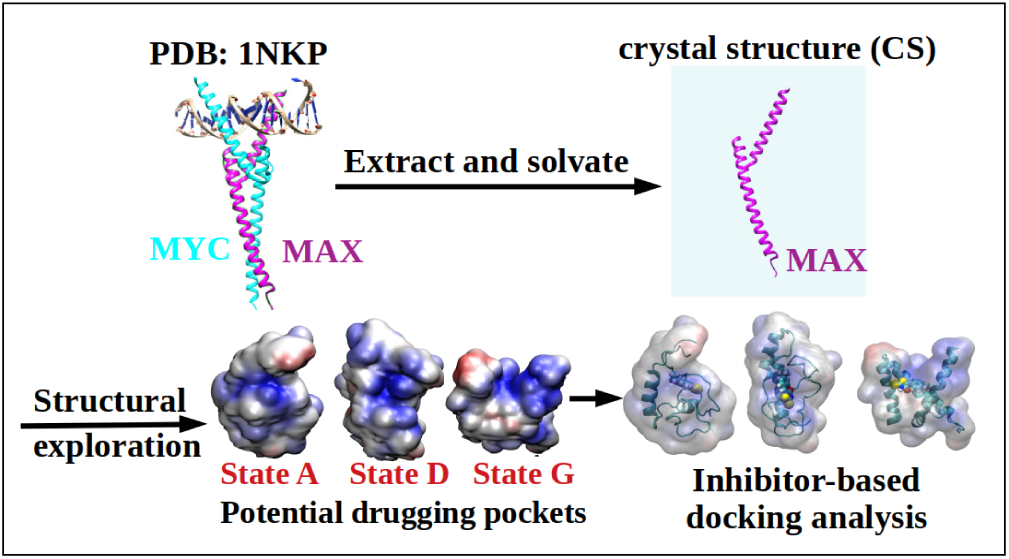

## Introduction

The V-myc avian myelocytomatosis viral oncogene homolog (MYC) and the MYC-Associated factor X (MAX) proteins are key transcription factors involved in regulating gene expression related to cell growth, proliferation, metabolism and apoptosis. MYC functions as a potent oncogene and requires dimerization with MAX to bind DNA at E-box sequences and activate transcription of target genes. MAX, in contrast, also forms dimers with other proteins like the MAX dimerization protein 1 (MXD1), which act as transcriptional repressors, thereby balancing MYC’s activity. The MYC-MAX network’s central role in tumorigenesis has become a significant target for anticancer therapies, although direct inhibition of MYC and/or MAX remains challenging due to its intrinsically disordered structure and widespread cellular functions.^1^ Whereas crystal structures of MAX-MYC dimers have been determined with precision, both including^2,3^ and excluding DNA,^4^ the determination of suitable drugging pockets is still quite elusive.

MAX encodes a ubiquitously expressed transcription factor that serves as an essential dimerization partner for MYC family proteins, facilitating their ability to regulate gene expression.^5,6^ MAX plays a pivotal role in modulating cellular processes such as proliferation, differentiation and apoptosis by forming heterodimers with MYC or other members of the MYC-MAX-MXD network.^7–9^ These interactions enable the binding of the resulting complexes to E-box sequences in DNA, where they activate or repress target genes depending on the dimeric partner involved. ^10,11^ Similarly to MYC, MAX possesses a basic helix-loop-helix leucine zipper (bHLHLZ) domain at its C-terminus, which is crucial for both dimerization and DNA binding.^12?^ However, in contrast to MYC, MAX lacks an intrinsic transactivation domain, acting instead as a scaffold that confers transcriptional activity or repression through its binding partners.^13,14^ The evolutionary conservation of MAX underscores its central role in balancing oncogenic and tumor-suppressive signals within the cell. ^15,16^

In many cancers, dysregulation of the MYC/MAX/MXD transcriptional network leads to an imbalance favoring MYC-MAX heterodimers, which drive uncontrolled cell proliferation and repress differentiation. Although MAX itself is not typically overexpressed or mutated, its obligatory role as a dimerization partner makes it a critical enabler of MYC-driven oncogenesis.^5,6^ MAX is expressed ubiquitously and forms stable bHLHLZ-mediated dimers with MYC, enabling binding to E-box DNA sequences and transcriptional activation of target genes involved in cell growth, metabolism and survival.^8,11^

Given the structural challenges in directly targeting MYC, an intrinsically disordered protein lacking well-defined binding pockets, MAX has emerged as an attractive alternative target for therapeutic intervention.^14?^ Disrupting the MYC-MAX interaction could effectively suppress MYC oncogenic activity without requiring direct inhibition of MYC itself. Several strategies have been proposed to this end, including the use of dominant-negative MYC mutants such as OmoMYC^17^ (OMO-103, currently in clinical trials), which sequesters MAX and blocks MYC-dependent transcription.^18^ Several small molecules, such as 10058-F4,^19^ KJ-Pyr-9,^20^ and MYCMI-6,^21^ have been developed to interfere with the MYC–MAX interaction directly. Also, peptides that mimic MAX or stabilize alternative dimerization partners (such as MXD proteins) could shift the transcriptional balance toward growth suppression.^7,9^

In order to identify viable drug-binding regions on MAX, it is essential to characterize its structural configurations. In this study, we leverage all-atom molecular dynamics (MD) simulations with high temporal and spatial resolution to model the conformational behavior of MAX and its potential interactions with MYC. We have chosen MD simulations due to its suitability to explore and characterize the conformational space of highly dynamic proteins.^22,23^ By analyzing the dynamic structural ensemble of MAX, we aim to uncover transient or stable pockets that could serve as potential drug-binding sites. Subsequently, we used the Well-Tempered Metadynamics (WT-MetaD)^24^ simulations to map the free energy landscape of MAX in solution, which will allow us to identify low-energy conformations with sufficient structural rigidity to accommodate small molecules, paving the way for the development of novel inhibitors targeting the MYC-MAX axis in cancer therapy.

## Methods

### System preparation and MD simulations

The aminoacid sequence of the MAX protein is available in UniProt (ID: P61244). There are several tridimensional structures of the MAX protein available in the Protein Data Bank, always in the form of a complex including another protein (MYC). We have selected the structure with PDB code 1NKP obtained in a detailed study of MAX recognition functions,^2^ but we should note that the MAX configuration is not different from that reported in other crystal structures. Hence, the MAX protein coordinates extracted from that PDB entry were employed as starting point for our simulations. Our simulation system consists of the MAX protein solvated in an explicit water box containing 0.15 M KCl to mimic the cell ionic strength. The system contains a total of 159,804 atoms. The system coordinates were generated using the CHARMM-GUI solution builder,^25–27^ and the CHARMM36m force field^28^ was adopted. All-atom MD simulations were conducted using AMBER24^29,30^ software. The system was minimized for 10000 steps followed by three 125 ps equilibrium runs while gradually reducing the harmonic constraints (NVT ensemble). Production runs were performed with the NPT ensemble, whereas Langevin dynamics with the friction coefficient 1.0 ps*^−^*^1^ was used for temperature regulation (310 K). The Monte Carlo barostat was used for the pressure regulation (1 bar). All bonds involving hydrogens were set to fixed lengths, allowing fluctuations of bond distances and angles for the remaining atoms. The time step was set to be 2 fs, and the frames were saved every 100 ps for analysis. The particle mesh Ewald method was automatically used by AMBER24 to calculate the electrostatic interactions, and the van der Waals interactions were calculated using a cutoff of 1.2 nm. Periodic boundary conditions were taken in three directions of space.

### Well-tempered metadynamics

The enhanced sampling method WT-MetaD applies a time-dependent biasing potential along a set of collective variables (CV) by adding a Gaussian additional bias potential to the total potential in order to overcome barriers larger than *k_B_T*, with *k_B_* being Boltzmann’s constant and *T* the temperature. In the method, sampling is performed on a selected number of degrees of freedom (*s_i_*), i.e. a chosen set of CV. For each degree of freedom, the biased potential *V* (*s_i_, t*) is a dynamical function constructed as the sum of Gaussian functions:

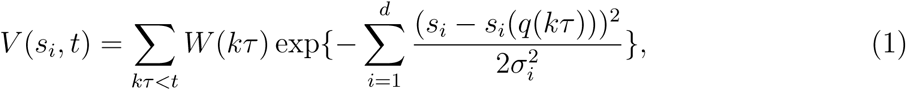

where *k* is an integer, *τ* is the Gaussian deposition stride, *W* (*kτ*) is the height of the Gaussian and *σ_i_* is the width of the Gaussian for the i-th CV. Full details are provided in Refs.^24,31,32^ In WT-MetaD, the biased potential can force the system to move around all possible states inside a particular range of the subspace of the selected CV. In the present work, we have taken a specific approach similar to previous works ^33–36^ where the CV were defined from distances between groups of particular amino acids. After analyzing the protein’s dynamics from the classical MD runs, we have selected two collective variables: *d1* and *d2*, where *d1* refers to the distance of center of mass of group1 (backbone atoms of residues 202-204) and group2 (backbone atoms of residues 282-284) and *d2* refers to the distance between C*_α_* atoms of Ile218 and Ile250 to explore the conformational space of MAX, see Section “Results and discussion” for a visualization of these CVs over the protein structures. The WT-MetaD simulations were performed using the PLUMED plugin (version 2.9.2) implemented in Gromacs MD package (version 2023.5).^37–39^ As initial configuration for the WT-MetaD simulations we considered the same relaxed crystal structure (CS) employed in the unbiased MD simulations. The main parameters of the WT-MetaD simulation are reported in Table 1. The V-rescale thermostat was considered for accurate temperature control and proper thermodynamic sampling at 310.15 K. Pressure was controlled to be 1 bar by a Parrinello-Rahman piston with a damping coefficient of 5 ps*^−^*^1^. Periodic boundary conditions in the three directions of space were also considered. More details, including analysis of the convergence of the metadynamics simulations (Figures S3-S7) can be found in the “Supporting Information” (SI).

**Table 1:** WT-MetaD simulation parameters used in this work.

| Parameter | Value |
| --- | --- |
| Gaussian width of $d1$ (nm) | 0.2 |
| Gaussian width of $d2$ (nm) | 0.2 |
| Gaussian hill (kJ/mol) | 1.2 |
| Deposition stride (ps) | 1 |
| Biasfactor | 8 |
| Simulation time (ns) | 3500 |

### Data analysis and visualization

The backbone atoms (N, C*_α_*, C, and O) of MAX were used to calculate the root mean squared deviation (RMSD) profile through RMSD Trajectory Tool of VMD-1.9.3.^40^ To explore the conformational landscape of the MAX protein during the molecular dynamics simulations, we carried out cluster analysis using the atomic coordinates of the backbone atoms, applying Principal Component Analysis (PCA)s.^41^ The analysis was performed using scikit-learn^42^ and MDAnalysis Python libraries.^43^ Pairwise distance matrices were constructed by extracting the positions of the backbone atoms throughout the entire trajectory using MDAnalysis. These structural descriptors were then employed as input data for PCA analysis using scikit-learn. Subsequently, the KMeans clustering algorithm^44^ was applied to the PCA-transformed data to partition the conformations into distinct clusters. Each cluster corresponds to a group of structurally similar protein conformations, potentially representing distinct metastable states sampled during the simulation. Then the representative structures from each cluster which are closest to the cluster centroid were identified for further structural characterization and visualization. This clustering approach provides a simplified yet informative view of the protein’s conformational ensemble, facilitating the interpretation of structural transitions and dynamic behavior. For each representative cluster structure, we report the angle between the two helices, calculated considering as the direction of each helix the one defined by the alpha carbon atoms of residues 210 and 221 of helix 1 and residues 249 and 267 of helix 2.

The R-package metadynminer^45,46^ was employed to analyze the WT-MetaD outputs. This package provides a set of tools for the efficient extraction, visualization, and interpretation of free energy surfaces (FES) computed from WT-MetaD trajectories. In this study, metadynminer was specifically employed to reconstruct the FES using the fast Bias Sum algorithm. Convergence of the free energy calculations was analyzed in detail in the Supporting Information. The *neb* function of the same package was employed to compute the minimum free energy paths (MFEP) by using the Nudged Elastic Band method. ^47^ MFEP describe the most probable transition routes between these states on the FES. The free energy profiles, free energy barriers and the transition states identified along the transitions were reported in detail in the main text and Supporting Information. The *neb* function of metadynminer also estimates the half-life of these states ^45^ using the classical Eyring–Polanyi equation of chemical kinetics,^48^ with a transmission coefficient assumed to be 1.

In addition to the full free energy map and the free energy profile along the MFEP, we have also computed the 1D free energy profiles F(*d*_1_) and F(*d*_2_) (where *d*_1_ and *d*_2_ are the CVs) by thermally averaging all contributions from *d*_2_ and *d*_1_ respectively in the full free energy profile F(*d*_1_,*d*_2_). More explicitly, F(*d*_1_) is calculated using:^49,50^

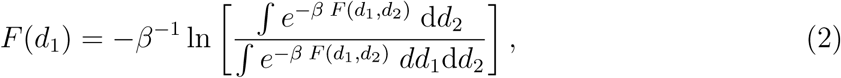

where *β* = 1*/*(*k*_B_*T*), *k*_B_ is the Boltzmann constant and T is the absolute temperature. *F* (*d*_2_) is obtained by simply exchanging the roles between *d*_1_ and *d*_2_ in Eq.(2). The analysis tool VMD^40^ and Chimera^51^ were used for visualization and analysis purposes.

## Results and discussion

### Structural transition pattern of MAX

The 3D structure of the MAX protein is shown in Fig. 1A, as obtained experimentally in a MYC-MAX dimer recognizing a common DNA target.^2^ The structure shows how the two helices of MAX are used to interact simultaneously with MYC and the DNA sequence in a way that has been intuitively compared with a specialized hand, perfectly designed to perform two tasks: shake hands with other proteins and grip onto DNA. Given this structure, a natural question is whether the MAX protein adopts a stable conformation when not dimerized with MYC—and, more broadly, what conformational states MAX can assume based on different arrangements of its two helices. This question should be relevant both for understanding how MAX performs its biological function but also should be relevant in the design of drugs interacting with MAX. To this end, we performed first exploratory MD simulations (described in this subsection) which were employed to identify relevant collective variables to be employed in subsequent extensive free energy calculations performed in the remaining subsections.

**Figure 1:**
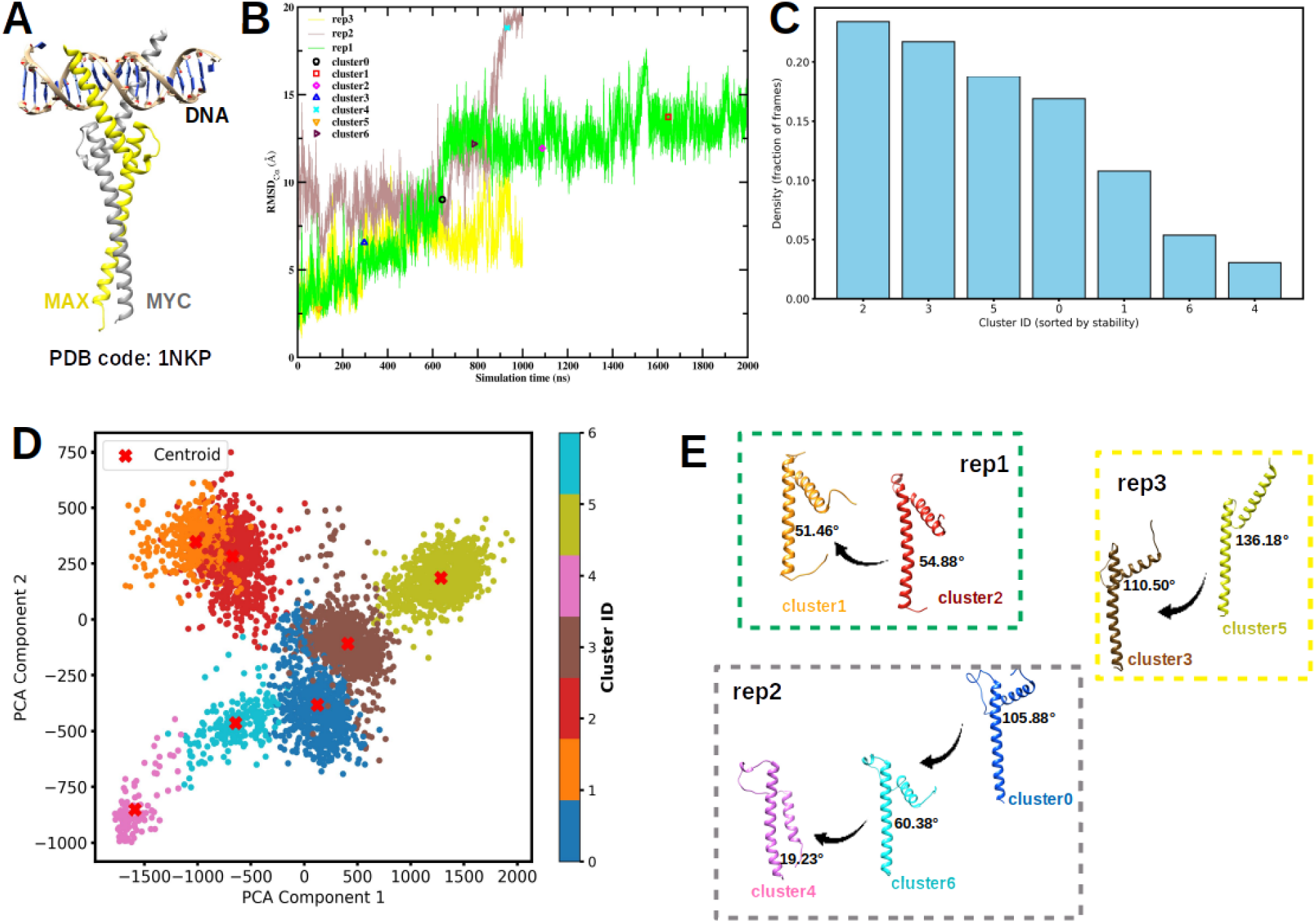
A. Crystal structure of a MYC-MAX dimer recognizing DNA obtained in Ref.^2^ B. RMSD profiles over the full simulation time of triplicate MD simulations initiated with different velocity seeds, spanning a total of 4 *µ*s. The frames corresponding to representative structures are highlighted. C: Cluster density profile derived from PCA clustering. D: Seven distinct clusters identified from PCA results, with the representative structure of each cluster indicated by a red cross. E. The transition pathways of MAX are illustrated as follows: from cluster 2 to cluster 1 in the first MD run (rep1, shown in subfigure B), from cluster 0 through cluster 6 to cluster 4 in the second independent MD run (rep2), and from cluster 5 to cluster 3 in the third MD run (rep3). The key structural transition is highlighted by the progressive decrease in the angle between the two helices.

We have performed triplicate MD simulations of the MAX protein solvated in salty water (150 mM of KCl mimicking physiological salt concentration), each initiated with distinct velocity seeds and spanning a total of 4 *µ*s. All simulations used the crystal structure of the MAX protein, extracted from the MYC–MAX–DNA complex (Fig. 1A) as the starting structure (see Section “Methods” for full details). During the MD trajectories the time evolution of quantities such as the RMSD (Fig. 1B) indicate the possibility of substantial changes in the protein conformation, even at the time scales explored by the unbiased MD simulations. In order to identify the essential features of the protein configurations explored during the MD trajectories, we have performed PCA analysis. We found that a 64.1% of the variance of the data can be explained by the first principal component. Adding a second component, the PC1 and PC2 variables explain 74.4% of the total variance (PC1 = 64.1%, PC2 = 10.3%). Adding more components does not significantly improve the variance explained, unless one adds a large number of PCA components (see Fig. S1 of SI). Therefore, we have decided to restrict our analysis to two PCA components. Once the two PCA components were obtained, we projected the trajectory into the PCA components and clustered the data (see Section “Methods”) and obtained representative structures for each cluster (Fig. 1C - 1E).

We evaluated clustering with varying numbers of clusters in order to identify the optimal choice. The results obtained with two different number of clusters (7 clusters and 8 clusters) are reported in Fig. 1 and Fig. S2, respectively. All these clusters correspond to a significant proportion of the trajectory configurations (Fig. 1C) and the structures characterizing these clusters are clearly different, as shown by the values of the PCA components (Fig. 1E), their different RMSD (Fig. 1B) and their different angle between the two helices that define the MAX protein structure (Fig. 1D). We found that 7 clusters were enough to represent the variability of the protein structures found in the MD simulation. Considering a larger number of clusters does not provide significantly different protein conformations, as illustrated with the results obtained for 8 clusters shown in the SI (Fig. S2). The results from the 8-cluster analysis were largely consistent with those obtained using 7 clusters, with the additional cluster further subdividing the existing groups without revealing fundamentally new structures. This consistency indicates that clustering into 7 groups captures the essential data structure and is sufficient for interpretation (Fig. 1).

Fig. 1C shows that cluster2 exhibits the highest population density, followed by cluster3, cluster5, cluster0, cluster1, cluster6, and cluster4 (note that the labeling of the clusters is arbitrary, not based on any physical quantity or time sequence). The time evolution between these clusters during the simulation was identified by tracking the simulation frames corresponding to each representative structure along the unbiased MD simulation timeline. Representative structures from the seven distinct clusters identified through PCA, along with the corresponding transition pathway, are shown in Fig. 1E. Across all three MD replicas, we observed a consistent structural transition in which the angle between the two helices in MAX progressively decreases along the trajectory. This reduction in inter-helical angle reflects a gradual compaction of the MAX domain, suggesting a shift toward a more closed conformational state. These results suggest that the observed transition is a reproducible and intrinsic feature of MAX’s conformational landscape and not an artifact of a particular simulation run.

### Selection of Collective Variables in WT-MetaD Simulations

After performing unbiased MD simulations, we will perform a more detailed analysis of the possible conformation space of MAX by employing WT-MetaD simulations, as described in the Section “Methods”. Before performing the simulations, the WT-MetaD technique requires the selection of the appropriate CV employed in the exploration of the conformation space. As it was mentioned above, we will consider as CV distances between particular aminoacids, as it was assumed in several previous works.^33,35?, 36^ The selection of these distances to be used as CV is made by analyzing the transitions between the seven different clusters obtained in the unbiased MD simulations (Fig. 1E). The geometrical changes undergone by the protein during these transitions can be clearly seen by aligning the representative structures of these clusters, as shown in Fig. 2A.

**Figure 2:**
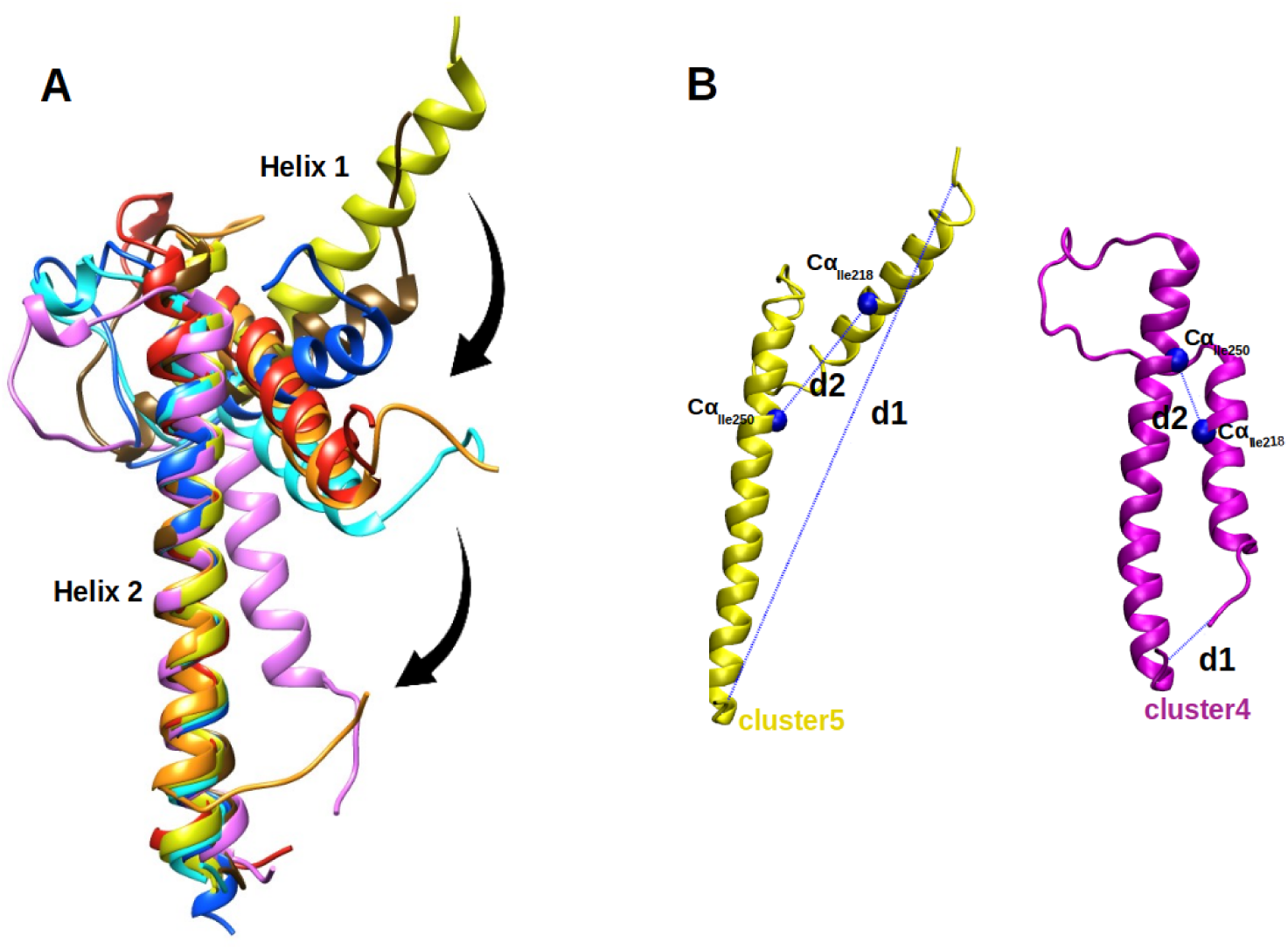
A. Representative structures from the seven clusters aligned based on residues 222-278 of Helix 2 of MAX, clearly illustrating the progression of conformational transitions. Colors correspond to those shown in Fig. 1E. B. Illustration of two collective variables (*d1* and *d2*), C*_α_* atoms of Ile218 and Ile250 are shown in Van der Waals (VdW) representation.

The conformational transitions are characterized by the rotation of helix 1 (residues 202–228) towards helix 2 (residues 239–281). This rotation ultimately leads to a significantly reduced angle between helix 1 and helix 2, as illustrated in Fig. 2A and a reduced distance between the N- and C-termini (indicated in Figure 2B as *d1*), along with a marked decrease in the distance between the C*_α_* atoms of residues Ile218 and Ile250 (indicated in Figure 2B as *d2*). As a result of this conformational rearrangement, the overall distance between the N- and C-termini of the 202–284 residue segment in unbound MAX is substantially reduced, leading to the formation of a more folded structure, represented by cluster4.

These observations were derived from the initial set of unbiased MD simulations conducted in the first phase of this study. Based on these structural changes, we selected *d1* and *d2* as the collective variables for subsequent enhanced sampling via WT-MetaD simulations.

### Free energy landscape of MAX

After selecting *d1* and *d2* as CV, we performed WT-MetaD simulations. The results are summarized in Fig. 3. We ensured that the free energy calculation is properly converged. As shown in Fig. 3, the free energy landscape of MAX reveals pronounced structural diversity within the low free energy region, populated by several metastable states. This indicates a rich ensemble of conformational states readily accessible in solution. The different states were found during the previous unbiased MD simulations. The results demonstrate that over the 4 *µ*s of an unbiased MD simulation, the system evolves from a high energy, crystal-like conformation of MAX (cluster5) towards a significantly lower-energy structure (cluster4). The free energy difference between the most stable state found in unbiased MD (cluster4) and the crystal-like state is ∼ 60 kJ/mol. Also, this figure shows that the unbiased MD simulations were not reaching the lowest free energy conformation of the protein, in spite of the long simulation time. These results emphasize that, even for a relatively small protein motif in this study (83 amino acids), the crystal structure can differ significantly from the minimum free energy configuration. They also underscore the importance of using enhanced sampling methods, such as WT-MetaD, to explore the structural variability of proteins. This enhanced sampling strategy enables us to explore the broader conformational landscape of MAX and ultimately identify previously inaccessible, stable drug-binding pockets in underexplored regions of the protein.

**Figure 3:**
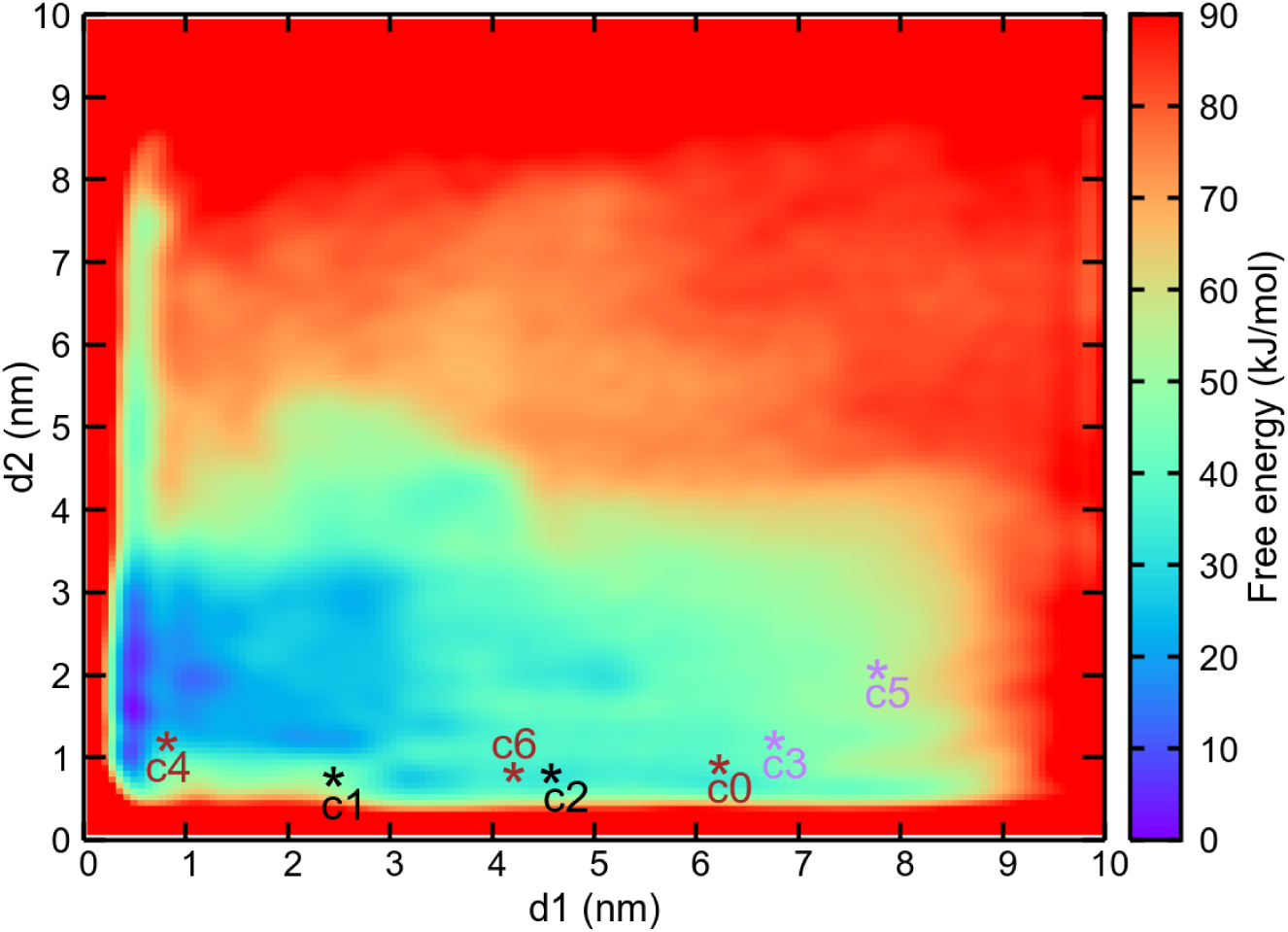
The two-dimensional free energy surface F(*d1*, *d2*) (kJ/mol) of unbound MAX in solution obtained from WT-MetaD simulations. Regions with free energy exceeding 90 kJ/mol are shown in red. ‘c0’ to ‘c6’ denote the coordinates of the representative structures from clusters 0 to 6 projected in this plot (see Fig. 1). Clusters c1 and c2 (from the rep1 MD simulation) are shown in black, c0, c4, and c6 (from rep2 MD simulation) are shown in brown, and c3 and c5 (from rep3 MD simulation) are shown in purple.

Looking in more detail at the free energy landscape, it is possible to observe several primary basins (Fig. 4). The details of these molecular structures (labeled A through L in Fig. 4) are also provided in Table 2.

**Figure 4:**
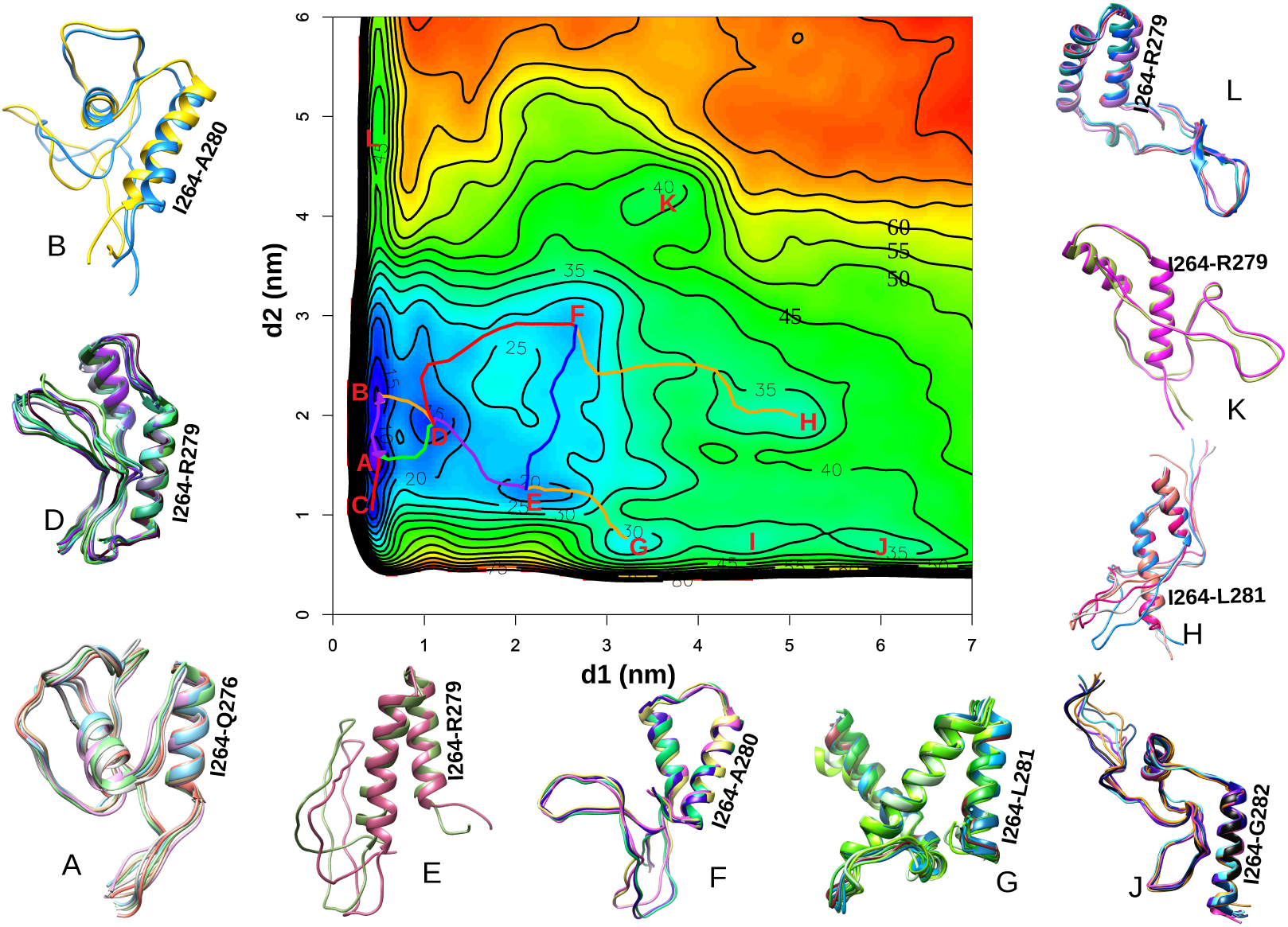
The partial free energy surface F(*d1*, *d2*) (kJ/mol) is presented here, highlighting newly discovered stable states within basins “A”–”L”. The corresponding MFEP connecting selected basins are also depicted. Representative MAX conformers from each basin are shown, highlighting the helical domain (I264-A280) near the C-terminus in all cases. Structures of MAX for basin C and I are not shown due to their short lifetime.

**Table 2:**
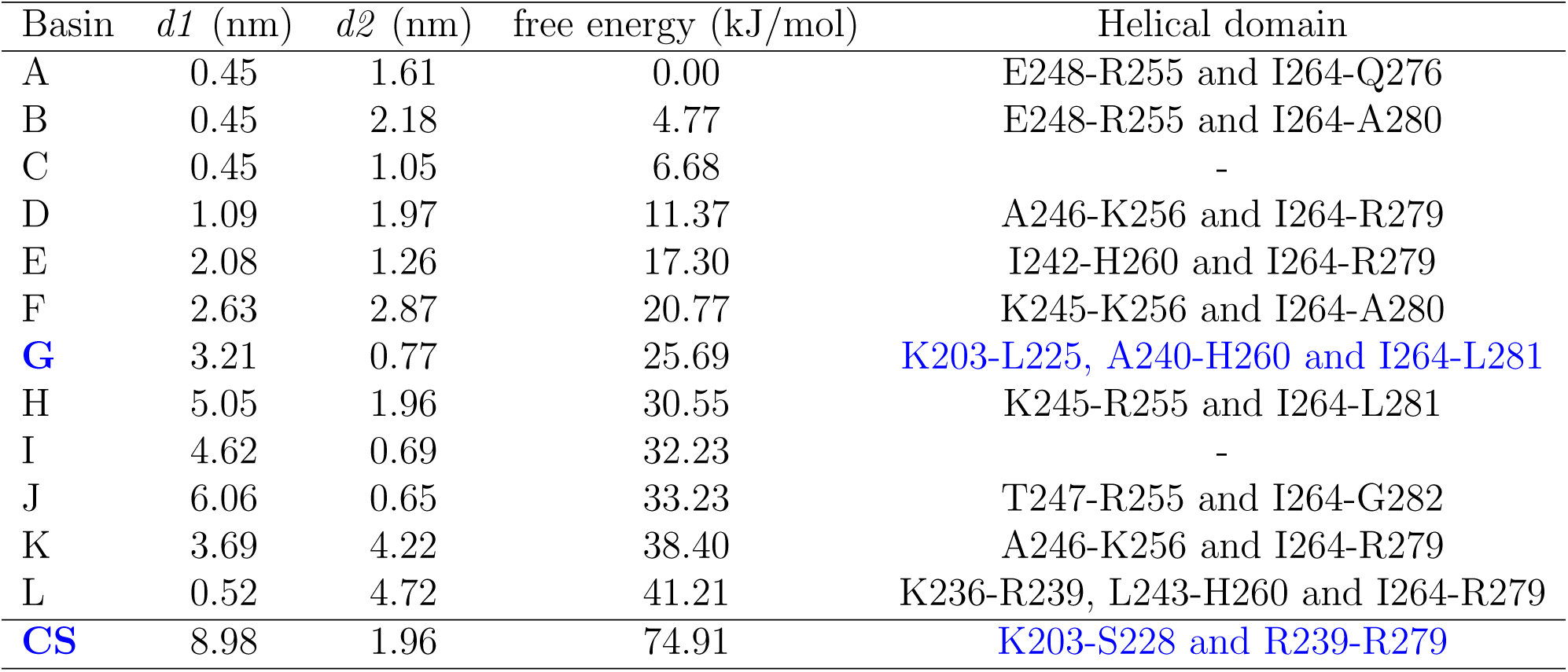
Coordinates of the representative basins (minima) corresponding to the Fig. 4 with the helical domain in each structure highlighted here. “CS” denotes the crystal structure in PDB 1NKP.

| Basin | $d1$ (nm) | $d2$ (nm) | free energy (kJ/mol) | Helical domain |
| --- | --- | --- | --- | --- |
| A | 0.45 | 1.61 | 0.00 | E248-R255 and I264-Q276 |
| B | 0.45 | 2.18 | 4.77 | E248-R255 and I264-A280 |
| C | 0.45 | 1.05 | 6.68 | - |
| D | 1.09 | 1.97 | 11.37 | A246-K256 and I264-R279 |
| E | 2.08 | 1.26 | 17.30 | I242-H260 and I264-R279 |
| F | 2.63 | 2.87 | 20.77 | K245-K256 and I264-A280 |
| **G** | 3.21 | 0.77 | 25.69 | K203-L225, A240-H260 and I264-L281 |
| H | 5.05 | 1.96 | 30.55 | K245-R255 and I264-L281 |
| I | 4.62 | 0.69 | 32.23 | - |
| J | 6.06 | 0.65 | 33.23 | T247-R255 and I264-G282 |
| K | 3.69 | 4.22 | 38.40 | A246-K256 and I264-R279 |
| L | 0.52 | 4.72 | 41.21 | K236-R239, L243-H260 and I264-R279 |
| **CS** | 8.98 | 1.96 | 74.91 | K203-S228 and R239-R279 |

Basins A, B, and C, located in the lower-left region of the free energy surface (*d1* ∼ 0.45 nm, *d2* ≤ 2.2 nm), represent the most thermodynamically stable conformations, as indicated by their deep free energy wells, see Table 2. These basins are tightly clustered, suggesting a compact ensemble of low-energy structures, associated with the native or folded state of MAX (see the corresponding structures of different basins in Fig. 4). Basins D, E, F, and G occupy moderately low-energy regions (11–26 kJ/mol) and may represent alternative metastable states that are structurally related to the native ensemble but differ in loop orientations or side-chain packing. In contrast, basins H-L, that are located in higher-energy regions (between 31–42 kJ/mol), refer to less stable and partially unfolded conformations. This free energy landscape reveals broader and more diffuse features in this region with greater structural flexibility. We observed a substantial free energy difference of 74.91 kJ/mol between the most stable basin (A) and the experimentally resolved crystal structure (CS). This substantial difference between the most stable configuration of MAX and its crystal structure has important implications for the development of drugs targeting MAX. This significant free energy gap underscores the difficulty of targeting MAX’s CS conformation with small molecules due to its low state population. The stable conformations identified in this study may offer promising targets for the development of effective inhibitors against MAX-associated diseases, much better than the crystal structure of MAX.

### Identification of transition paths and transition states

The WT-MetaD technique reveals not only the free energy basins (the most stable configurations) but also the free energy barriers that the protein needs to overcome to shift between stable states. Furthermore, the maxima along each path give indications of the approximated location of transition states (TS) between stable states. In table 3 we show the free energy barriers between the different basins of interest seen in Fig. 4. The transition half-lives between these basins, estimated using the Eyring–Polanyi equation (see Section “Methods”), are also provided. The corresponding free energy profiles along the MFEP for each case are shown in Fig. S8 of SI, including the estimated locations of local TS between stable basins, and the raw data can be found in our Github repository.^52^ The energy barriers between low-energy basins and higher-energy regions (e.g., A to D, E to G, and F to H) suggest that large-scale rearrangements are thermodynamically unfavorable and likely require crossing high activation barriers. The half-life values along the MFEP between basins are consistent with the number of conformers sampled for each basin in the WT-MetaD trajectories. A shorter half-life indicates a faster kinetic transition between states for MAX, leading to lower accumulation in individual state populations (i.e. fewer structurally relevant conformers). Notably, the structures of MAX corresponding to basin C and I, which exhibit very short transition half-lives (0.32 ps and 0.27 ps, respectively), were not captured in the WT-MetaD output, where frames were saved every 2 ps, indicating that they are transient states.

**Table 3:**
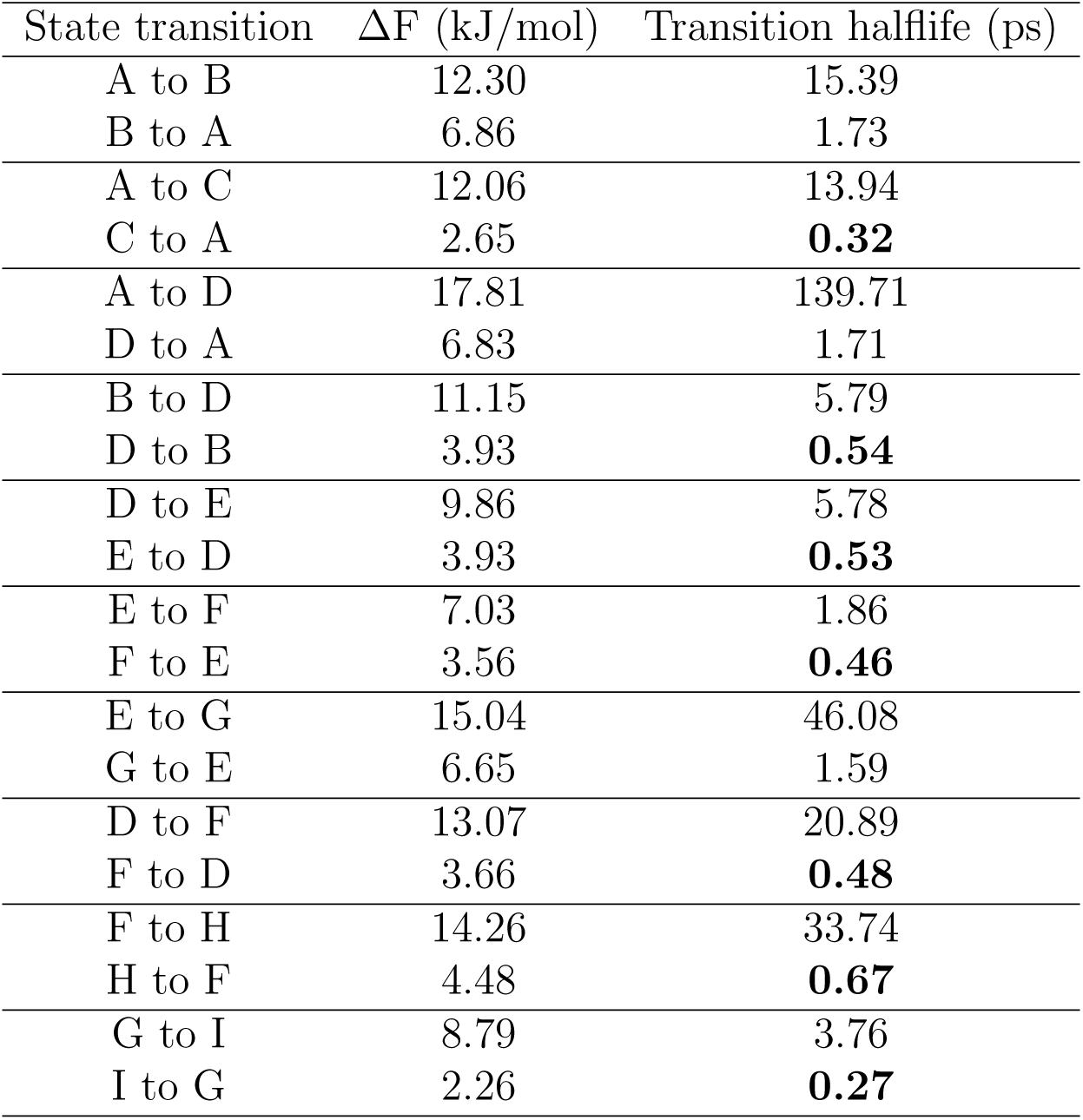
free energy barriers (ΔF) between basins (kJ/mol) along the corresponding MFEP and their transition half-lives (ps), as predicted by the Nudged Elastic Band method and Eyring–Polanyi equation, respectively (see Section “Methods”). State transitions with half-lives shorter than 1 ps are shown in bold characters.

In addition to the complex picture provided by the MFEP, it is possible to obtain a more simplified but useful picture by considering one-dimensional (1D) profiles where free energy depends of only one CV, after the other CV has been integrated out over their Boltzmann factor. Previous works^49^ have shown the relevance of these 1D free energy profiles since, under appropriate conditions, they can be directly related to experimentally measured free energies Δ*G*. In Fig. 5 we show the results for the 1D free energy profiles F(*d*_1_) and F(*d*_2_) obtained by thermally averaging all contributions from *d*_2_ and *d*_1_ respectively (see the Section “Methods” for further details).

**Figure 5:**
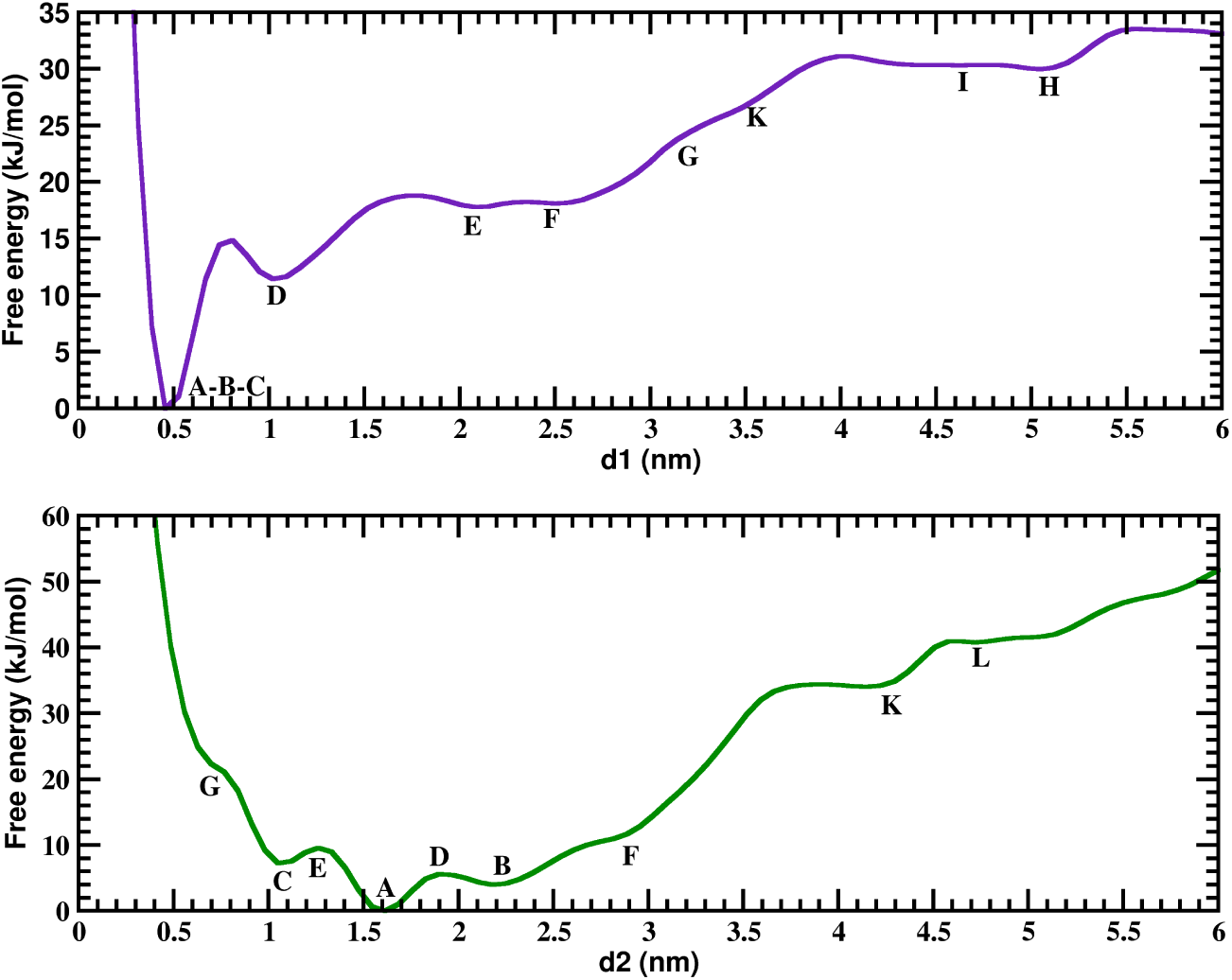
1D integrated free energy profiles for two CVs at 3500 ns. Basin regions marked with same labels (A-L) as in Fig. 4. In order to directly compute the height of free energy barriers, absolute minima of each free energy are set equal to zero.

The obtained free energy profiles F(*d*_1_) and F(*d*_2_) are rather smooth, with less features than the MFEP free energy profiles shown in Fig. S8. The only remarkable feature of these profiles is seen in the case of F(*d*_1_) in the region connecting basins A and D. It should be noted that this is the transition which involves the largest free energy and largest transition half-life, as identified by the MFEP (see Table 3). The F(*d*_1_) profile between A and D states presents a small local maxima close to D and a deep energy minimum around A. Interestingly, the profile of *F* (*d*_1_) in this region (Fig. 5) is close to the MFEP profile between these same states (Fig. S8). The free energy difference involved in the transition between A and D following the MFEP path is 17.8 kJ/mol (see Table 3) and the prediction from of *F* (*d*_1_) is about 15 kJ/mol (Fig. 5). This comparison demonstrates that although the transition between A and D involves changes in both CV (see Fig. 4), the free energy changes are essentially due to only one of the CV (in this case *d*_1_). This is not the case for most of the other transitions shown in Table 3, as seen by comparing Fig. 5 with Fig. S8, which involve more complex free energy contributions from changes in both CV and that need to be described by the MFEP connecting them.

Overall, the results for the free energy barriers found in the calculations reported here show many possible transitions between free energy basins involving free energies much larger than the thermal energy at 310K (∼ 2.6 kJ/mol). They can be compared to other biological bottlenecks, such as those related to the RAS family of oncogenes, where barriers of ∼ 20 kJ/mol due to changes in specific distances and orientations were estimated for the activation of mutated RAS proteins by Fourier Transform Infrared Spectroscopy^53^ and simulations^54^ or to the case of KRAS-G12D, when bound to a cell membrane, where angular barriers in the range 8-17 kJ/mol were estimated.^55^

### Location of potential targeting pockets at the surface of MAX

After characterizing the possible configurations of MAX, our interest now is to identify which of these configurations are of interest for *in silico* drug design efforts. A possible therapeutic strategy could be to target unbound MAX, limiting its availability for MYC binding, as described in the Section “Introduction”. We should note here that the crystal structure employed as initial condition in our simulations corresponds to a MAX protein in a MYC-MAX dimer. The unbound MAX protein in solution can be expected to be in one of the free energy basins described in the previous subsections. Therefore, to identify druggable targets from MAX configurations we should explore the configurations and free energy landscape connecting the minimum free energy state identified in our calculations (state A in Table 2) with the CS structure.

By following the MFEP from the most stable state (basin A) to the CS, it is clear that the MFEP, depicted in red in Fig. 6A, passes through basin D, leading to a high free energy barrier of approximately 75 kJ/mol. Interestingly, structure G shows a high degree of similarity to the CS within the helical domains when compared to other basins (highlighted in blue in Table 2). Upon aligning the structures of basin G and CS, it becomes evident that MAX only needs to rotate residues 202–225 by 164.2° and residues 264–284 by 133.1° to adopt the conformation of CS (see Fig. 6B). This suggests that state G may represent a transition state to adopt the conformation of the CS during the process of the MAX-MAX/MAX-MYC dimerization. Given the higher population of state G, reflected by the greater number of conformers compared to other basins in Fig. 4, we further examined the MFEP from basin A to G, which proceeds through basin D and involves a free energy barrier of 32.01 kJ/mol. Additionally, the MFEP from basin G to the CS revealed a free energy difference of 49.60 kJ/mol, which is smaller than that from basin A to the CS. These findings highlight states A as the most stable structure of MAX in solution, D and G as key intermediates, suggesting their potential as promising therapeutic targets in MAX-associated diseases. An interesting feature of these structures is the fact that they expose positively charged regions. In Fig. 6C we show the electrostatic potential of the representative structures of the A, D and G states calculated at the Poisson-Boltzmann level of theory using the APBS^56^ software. As seen in that figure, there are large pockets of these structures at potentials around ∼ 80 mV. This suggests electrostatic interaction as a suitable mechanism for binding to MAX. Fig. 6D shows the charge distribution of the small-molecule c-MYC inhibitor 10058-F4, which has been proposed to interact with both MAX and MYC and inhibit their heterodimerization.^19^

**Figure 6:**
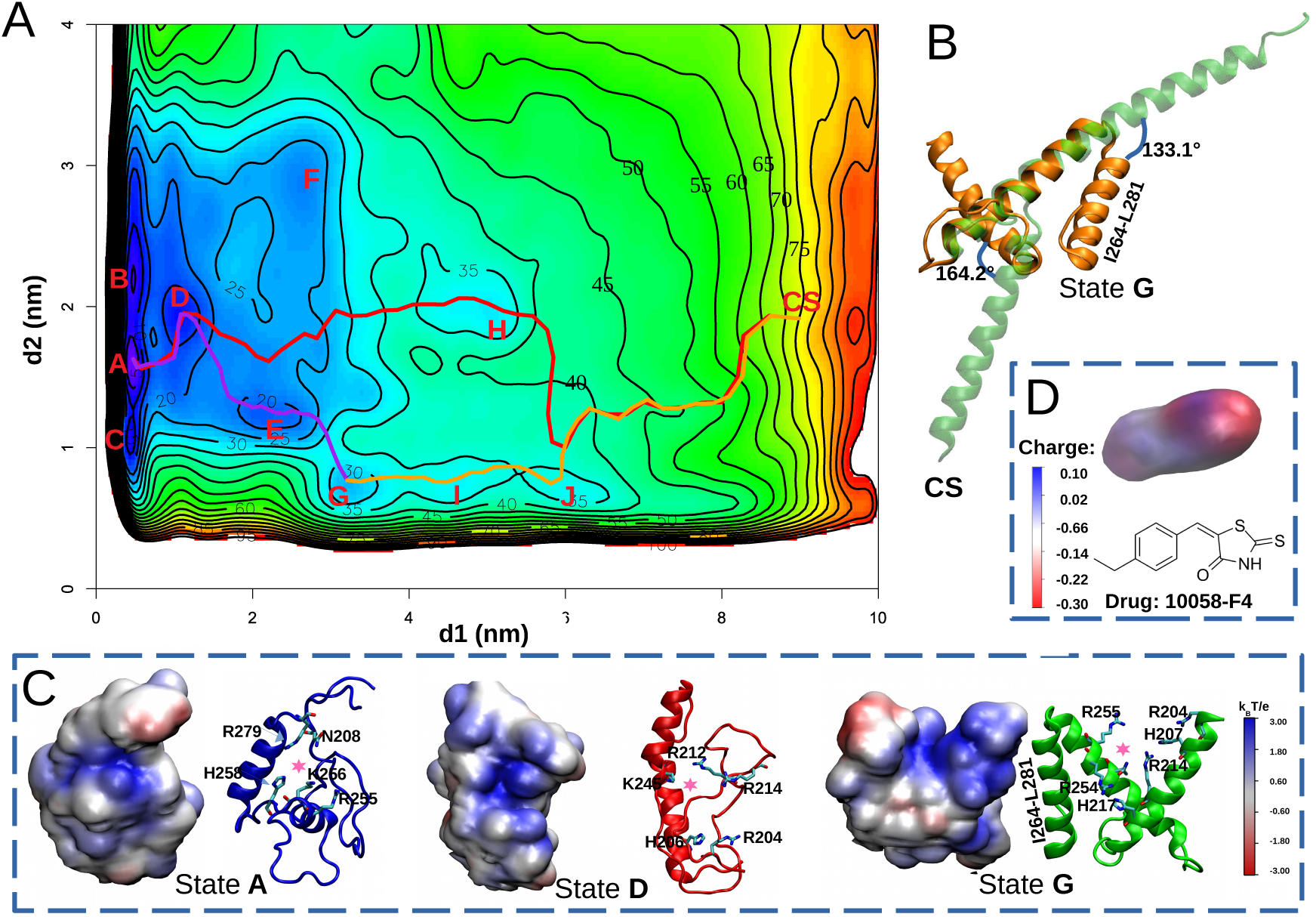
A: MFEPs between A to CS (red), A to G (purple), and G to CS (orange) on the free energy surface F(*d1*, *d2*) (kJ/mol). B: Aligned structures of state G and the CS. Angles between vectors along the helical domain are shown here. C: Electrostatic potential of MAX in states A, D, and G in units of *k_B_T/e* (*k_B_T/e* ≈ 26.7 mV). The potential drug-binding pocket of each state is marked with a pink star, and positive side chains of these pockets are highlighted here. D: Charge distribution of small-molecule c-MYC inhibitor 10058-F4.

Our results suggest that the binding of this molecule to states A, D, and G of MAX should be possible, driven by electrostatic complementarity. To verify this hypothesis, we conducted additional docking analyses between MAX and the 10058-F4 inhibitor. Full details are reported in the Supporting Information. The docking outcomes, presented in Fig. 7, show the most favorable complex structures for states A, D, and G and align well with our preliminary predictions in Fig. 6C. Coordinates of these complexes are available in our GitHub repository.

**Figure 7:**
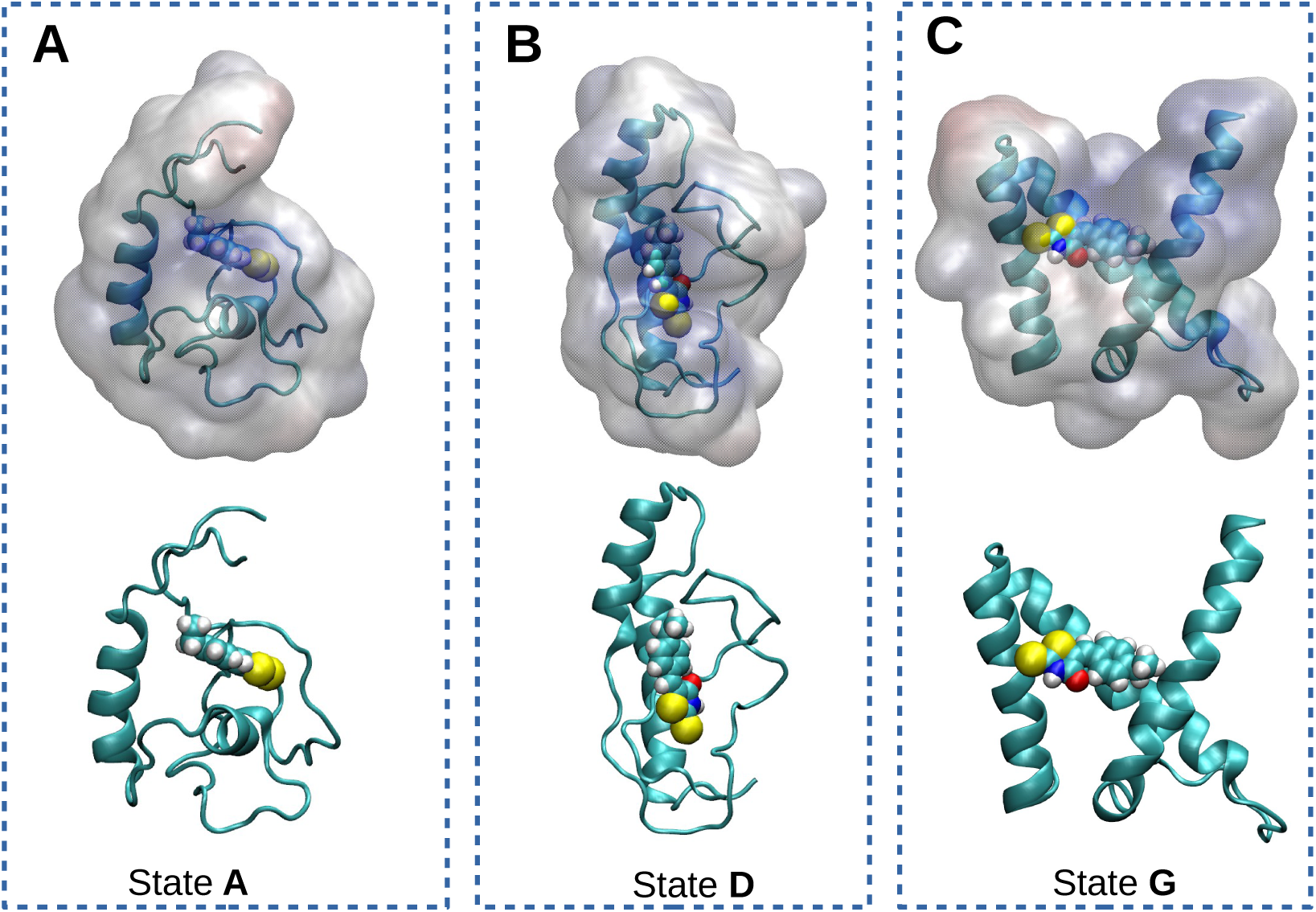
The most favorable structures of MAX-inhibitor complex predicted by HAD-DOCK2.4.^57^ The inhibitor 10058-F4 is displayed in van der Waals representation in all cases. Figures were generated using VMD.

## Conclusions

The importance of the MYC–MAX dimer lies in its ability to associate with DNA in the cell nucleus and, more critically, in the essential role this complex plays in the dysregulation of gene expression in roughly 70% of all known cancers. Despite extensive research, no effective strategies for directly targeting MYC have yet been developed. Because direct inhibition of MYC has proven extremely challenging, MAX has emerged as an attractive alternative target for novel therapeutic approaches.^14?^ In this context, the present work reports a detailed study of the FES of MAX in solution, aiming to identify previously uncharacterized druggable pockets associated with the most relevant minima of the FES. We have observed that the monomeric, unbound MAX motif exhibits a rich conformational ensemble, as revealed by extensive classical MD (2 *µ*s) and WT-MetaD simulations (3.5 *µ*s). Classical MD runs revealed five dominant structural clusters, which guided the selection of two distances as CV for the subsequent WT-MetaD enhanced sampling to further explore the structural diversity of MAX.

The 2D FES constructed from WT-MetaD simulations revealed a broad conformational space, featuring numerous low-energy basins corresponding to metastable states. The most stable basin (A) lies ∼75 kJ/mol below the experimentally resolved crystal structure, indicating that the latter is a low-populated state in solution. Basins A and B form a compact cluster of highly stable conformations with tightly packed helices, while D–G represent moderately stable states. By contrast, higher-energy basins (H–L) are associated with partially unfolded or extended conformations, underscoring the intrinsic flexibility of MAX. The Minimum free energy Path analysis using the package R-Metadynminer indicated that the transition from A to D exhibits the largest energy barrier (Δ F ∼ 18 kJ/mol), suggesting a significant kinetic bottleneck. In contrast, transitions among other basins (e.g., A to B or B to D) have lower energy barriers and shorter lifetimes. Certain basins such as C and I were identified as highly transient (lifetime ≤ 0.5 ps), leading to their absence in the WT-MetaD output frames. This indicates that not all minima contribute equally to the long-lived conformational ensemble of MAX.

Importantly, the substantial energy gap between the stable solution-phase conformations and the crystal structure highlights the need to target MAX in its native, lowest-energy states. The stable conformations identified in the present study may therefore represent promising candidates for the development of effective inhibitors, extending beyond the long-studied crystal structure of the MYC–MAX complex. In particular, the identification of persistent pockets within low-energy basins (particularly A, D, and G) provides promising templates for rational structure-based drug design against the MYC–MAX interaction interface. Furthermore, electrostatic potential maps of states A, D, and G reveal positively charged pockets, while the small-molecule MYC inhibitor 10058-F4 exhibits a negatively charged electrostatic profile, suggesting potential electrostatic complementarity that could be exploited for binding. The resulting complex structures predicted by additional docking analyses between MAX and the 10058-F4 inhibitor align well with our predictions.

## Data and Software Availability

The raw data and analysis scripts that support the findings of this study can be found in the corresponding repository of Github: https://github.com/HuixiaLuScienceRocks/Probing_the_Structural_Dynamics_of_Unbound_MAX_Protein_Insights_from_Well_Tempered_Metadynamics. The repository contains input files for WT-MetaD simulations, coordinates of meta- and stable MAX structures, scripts for PCA and angle calculations, raw block analysis data, minimum free energy pathways between different basins, optimized structure of 10058-F4, along with docking results for 10058-F4 and the most favorable MAX–inhibitor complex structures predicted through docking simulations.^52^

## Supporting Information

Supplementary figures and further information: outcomes of the PCA analysis; convergence assessments for the WT-MetaD simulations; free energy profiles along the Minimum Free Energy Paths connecting selected basins; computational details for the DFT-based structural optimization of the inhibitor 10058-F4; supplemental docking methodology.

## Author Contributions

Huixia Lu: Writing – original draft, Writing – review & editing, Visualization, Validation, Software, Methodology, Investigation, Formal analysis, Conceptualization.

Jordi Marti: Writing – original draft, Writing – review & editing, Resources, Conceptualization, Funding acquisition.

Jordi Faraudo: Writing – original draft, Writing – review & editing, Resources, Conceptualization, Funding acquisition.

## Acknowledgments

This work was supported by grants PID2021-124297NB-C32, PID2021-124297NB-C33, PID-2024-157478NB-C32 and PID2024-157478NB-C33 funded by MCIN/AEI/10.13-039/5011000-11033 and the “Severo Ochoa” grant CEX2023-001263-S for Centers of Excellence awarded to ICMAB-CSIC. J.M. thanks the *Generalitat de Catalunya* for the support through the grant *Grup de Recerca SGR-Cat2021 Condensed, Complex and Quantum Matter Group* reference 2021SGR-01411 and to the Polytechnic University of Catalonia-Barcelona Tech through the funding AGRUPS. The authors thankfully acknowledge computreNos-trum5 supercomputer as well as technical support provided by BSC under project number RES-BCV-2025-1-0008.

## References

(1) Schütz, S.; Bergsdorf, C.; Goretzki, B.; Lingel, A.; Renatus, M.; Gossert, A. D.; Jahnke, W. The disordered MAX N-terminus modulates DNA binding of the transcription factor MYC: MAX. Journal of molecular biology 2022, 434, 167833.

(2) Nair, S. K.; Burley, S. K. X-ray structures of Myc-Max and Mad-Max recognizing DNA: molecular bases of regulation by proto-oncogenic transcription factors. Cell 2003, 112, 193–205.

(3) Beaulieu, M.-E.; Castillo, F.; Soucek, L. Structural and biophysical insights into the function of the intrinsically disordered Myc oncoprotein. Cells 2020, 9, 1038.

(4) Sammak, S.; Hamdani, N.; Gorrec, F.; Allen, M. D.; Freund, S. M.; Bycroft, M.; Zinzalla, G. Crystal structures and nuclear magnetic resonance studies of the apo form of the c-MYC: MAX bHLHZip complex reveal a helical basic region in the absence of DNA. Biochemistry 2019, 58, 3144–3154.

(5) Blackwood, E. M.; Eisenman, R. N. Max: A Helix-Loop-Helix Zipper Protein That Forms a Sequence-Specific DNA-Binding Complex with Myc. Science 1991, 251, 1211–1217.

(6) Grandori, C.; Cowley, S. M.; James, L. P.; Eisenman, R. N. The Myc/Max/Mad network and the transcriptional control of cell behavior. Annul Review of Cell and Developmental Biology 2000, 16, 653–699.

(7) Mateyak, M. K.; Obaya, A. J.; Adachi, S.; Sedivy, J. M. Phenotypes of c-Myc-deficient rat fibroblasts isolated by targeted homologous recombination. Cell Growth Differentiation 1997, 8, 1039–1048.

(8) Lüscher, B. Function and regulation of the transcription factors of the Myc/Max/Mad network. Gene 2001, 277, 1–14.

(9) Sabo, A.; Kress, T. R.; Pelizzola, M.; others Selective transcriptional regulation by Myc in cellular growth control and lymphomagenesis. Nature 2014, 511, 488–492.

(10) Lüscher, B. Function and regulation of the transcription factors of the Myc/Max/Mad network. Gene 2001, 277, 1–14.

(11) Allevato, M.; Bolotin, E.; Grossman, M.; Mane-Padros, D.; Sladek, F. M.; Martinez, E. Sequence-specific DNA binding by MYC/MAX to low-affinity non-E-box motifs. PloS one 2017, 12, e0180147.

(12) Cascón, A.; Robledo, M. MAX and MYC: a heritable breakup. Cancer research 2012, 72, 3119–3124.

(13) Kretzner, L.; Blackwood, E. M.; Eisenman, R. N. Myc and Max proteins possess distinct transcriptional activities. Nature 1992, 359, 426–429.

(14) Meyer, N.; Penn, L. Z. Reflecting on 25 years with MYC. Nature Reviews Cancer 2008, 8, 976–990.

(15) Augert, A.; Mathsyaraja, H.; Ibrahim, A. H.; Freie, B.; Geuenich, M. J.; Cheng, P.-F.; Alibeckoff, S. P.; Wu, N.; Hiatt, J. B.; Basom, R.; others MAX functions as a tumor suppressor and rewires metabolism in small cell lung cancer. Cancer cell 2020, 38, 97–114.

(16) Young, S. L.; Diolaiti, D.; Conacci-Sorrell, M.; Ruiz-Trillo, I.; Eisenman, R. N.; King, N. Premetazoan ancestry of the Myc–Max network. Molecular biology and evolution 2011, 28, 2961–2971.

(17) Garralda, E.; Beaulieu, M.-E.; Moreno, V.; Casacuberta-Serra, S.; Martinez-Martin, S.; Foradada, L.; Alonso, G.; Masso-Valles, D.; López-Estévez, S.; Jauset, T.; others MYC targeting by OMO-103 in solid tumors: a phase 1 trial. Nature medicine 2024, 30, 762–771.

(18) Soucek, L.; Jucker, R.; Panacchia, L.; Ricordy, R.; Tato, F.; Nasi, S. Omomyc, a potential Myc dominant negative, enhances Myc-induced apoptosis. Cancer research 2002, 62, 3507–3510.

(19) Follis, A. V.; Hammoudeh, D. I.; Daab, A. T.; Metallo, S. J. Small-molecule pertubation of competing interactions between c-Myc and Max. Bioorganic & medicinal chemistry letters 2009, 19, 807–810.

(20) Hart, J. R.; Garner, A. L.; Yu, J.; Ito, Y.; Sun, M.; Ueno, L.; Rhee, J.-K.; Baksh, M. M.; Stefan, E.; Hartl, M.; others Inhibitor of MYC identified in a Kröhnke pyridine library. Proceedings of the National Academy of Sciences 2014, 111, 12556–12561.

(21) AlSultan, D.; Kavanagh, E.; O’Grady, S.; Eustace, A. J.; Castell, A.; Larsson, L.-G.; Crown, J.; Madden, S. F.; Duffy, M. J. The novel low molecular weight MYC antagonist MYCMI-6 inhibits proliferation and induces apoptosis in breast cancer cells. Investigational new drugs 2021, 39, 587–594.

(22) Lindorff-Larsen, K.; Trbovic, N.; Maragakis, P.; Piana, S.; Shaw, D. E. Structure and Dynamics of an Unfolded Protein Examined by Molecular Dynamics Simulation. Journal of the American Chemical Society 2012, 134, 3787–3791.

(23) Larsen, A. H.; Wang, Y.; Bottaro, S.; Grudinin, S.; Arleth, L.; Lindorff-Larsen, K. Combining molecular dynamics simulations with small-angle X-ray and neutron scattering data to study multi-domain proteins in solution. PLoS computational biology 2020, 16, e1007870.

(24) Barducci, A.; Bussi, G.; Parrinello, M. Well-tempered metadynamics: a smoothly converging and tunable free-energy method. Physical review letters 2008, 100, 020603.

(25) Jo, S.; Kim, T.; Iyer, V. G.; Im, W. CHARMM-GUI: a web-based graphical user interface for CHARMM. Journal of computational chemistry 2008, 29, 1859–1865.

(26) Brooks, B. R.; Brooks III, C. L.; Mackerell Jr, A. D.; Nilsson, L.; Petrella, R. J.; Roux, B.; Won, Y.; Archontis, G.; Bartels, C.; Boresch, S.; others CHARMM: the biomolecular simulation program. Journal of computational chemistry 2009, 30, 1545–1614.

(27) Lee, J.; Cheng, X.; Jo, S.; MacKerell, A. D.; Klauda, J. B.; Im, W. CHARMM-GUI input generator for NAMD, GROMACS, AMBER, OpenMM, and CHARMM/OpenMM simulations using the CHARMM36 additive force field. Biophysical journal 2016, 110, 641a.

(28) Huang, J.; MacKerell Jr, A. D. CHARMM36 all-atom additive protein force field: Validation based on comparison to NMR data. Journal of computational chemistry 2013, 34, 2135–2145.

(29) Case, D. A.; Cheatham III, T. E.; Darden, T.; Gohlke, H.; Luo, R.; Merz Jr, K. M.; Onufriev, A.; Simmerling, C.; Wang, B.; Woods, R. J. The Amber biomolecular simulation programs. Journal of computational chemistry 2005, 26, 1668–1688.

(30) Case, D. A.; Aktulga, H. M.; Belfon, K.; Cerutti, D. S.; Cisneros, G. A.; Cruzeiro, V. W. D.; Forouzesh, N.; Giese, T. J.; Gotz, A. W.; Gohlke, H.; others AmberTools. Journal of chemical information and modeling 2023, 63, 6183–6191.

(31) Bussi, G.; Laio, A.; Parrinello, M. Equilibrium free energies from nonequilibrium meta-dynamics. Physical review letters 2006, 96, 090601.

(32) Bonomi, M.; Camilloni, C.; Vendruscolo, M. Metadynamic metainference: Enhanced sampling of the metainference ensemble using metadynamics. Scientific reports 2016, 6, 31232.

(33) Lovera, S.; Morando, M.; Pucheta-Martinez, E.; Martinez-Torrecuadrada, J. L.; Saladino, G.; Gervasio, F. L. Towards a molecular understanding of the link between imatinib resistance and kinase conformational dynamics. PLoS computational biology 2015, 11, e1004578.

(34) Lu, H.; Marti, J. Cellular absorption of small molecules: free energy landscapes of melatonin binding at phospholipid membranes. Scientific reports 2020, 10, 9235.

(35) Lu, H.; Martí, J. Influence of cholesterol on the orientation of the farnesylated GTP-bound KRas-4B binding with anionic model membranes. Membranes 2020, 10, 364.

(36) Singh, A.; Lakkaniga, N. R. Exploring the Conformational Space of MPS1 Kinase Using Metadynamics. Proteins: Structure, Function, and Bioinformatics 2025, 93, 1118–1127.

(37) Bonomi, M.; Branduardi, D.; Bussi, G.; Camilloni, C.; Provasi, D.; Raiteri, P.; Donadio, D.; Marinelli, F.; Pietrucci, F.; Broglia, R. A.; others PLUMED: A portable plugin for free-energy calculations with molecular dynamics. Computer Physics Communications 2009, 180, 1961–1972.

(38) Tribello, G. A.; Bonomi, M.; Branduardi, D.; Camilloni, C.; Bussi, G. PLUMED 2: New feathers for an old bird. Computer physics communications 2014, 185, 604–613.

(39) Promoting transparency and reproducibility in enhanced molecular simulations. Nature methods 2019, 16, 670–673.

(40) Humphrey, W.; Dalke, A.; Schulten, K. VMD: visual molecular dynamics. Journal of molecular graphics 1996, 14, 33–38.

(41) Greenacre, M.; Groenen, P. J.; Hastie, T.; d’Enza, A. I.; Markos, A.; Tuzhilina, E. Principal component analysis. Nature Reviews Methods Primers 2022, 2, 100.

(42) Pedregosa, F.; Varoquaux, G.; Gramfort, A.; Michel, V.; Thirion, B.; Grisel, O.; Blondel, M.; Prettenhofer, P.; Weiss, R.; Dubourg, V.; others Scikit-learn: Machine learning in Python. the Journal of machine Learning research 2011, 12, 2825–2830.

(43) Gowers, R. J.; Linke, M.; Barnoud, J.; Reddy, T. J. E.; Melo, M. N.; Seyler, S. L.; Domanski, J.; Dotson, D. L.; Buchoux, S.; Kenney, I. M.; others MDAnalysis: a Python package for the rapid analysis of molecular dynamics simulations; 2019.

(44) Likas, A.; Vlassis, N.; Verbeek, J. J. The global k-means clustering algorithm. Pattern recognition 2003, 36, 451–461.

(45) Trapl, D.; Spiwok, V. Analysis of the Results of Metadynamics Simulations by metadynminer and metadynminer3d. arXiv preprint arXiv:2009.02241 2020,

(46) Hošek, P.; Spiwok, V. Metadyn View: Fast web-based viewer of free energy surfaces calculated by metadynamics. Computer Physics Communications 2016, 198, 222–229.

(47) Henkelman, G.; Jónsson, H. Improved tangent estimate in the nudged elastic band method for finding minimum energy paths and saddle points. The Journal of chemical physics 2000, 113, 9978–9985.

(48) Eyring, H. The Activated Complex in Chemical Reactions. The Journal of Chemical Physics 1935, 3, 107–115

(49) Jambeck, J. P.; Lyubartsev, A. P. Exploring the free energy landscape of solutes embedded in lipid bilayers. The Journal of Physical Chemistry Letters 2013, 4, 1781–1787.

(50) Roux, B. Statistical mechanical equilibrium theory of selective ion channels. Biophysical journal 1999, 77, 139–153.

(51) Pettersen, E. F.; Goddard, T. D.; Huang, C. C.; Couch, G. S.; Greenblatt, D. M.; Meng, E. C.; Ferrin, T. E. UCSF Chimera—a visualization system for exploratory research and analysis. Journal of computational chemistry 2004, 25, 1605–1612.

(52) Lu, H. Project MAX. https://github.com/HuixiaLuScienceRocks/Probing_the_Structural_Dynamics_of_Unbound_MAX_Protein_Insights_from_Well_Tempered_Metadynamics, 2025.

(53) Li, Y.; Zhang, Y.; Großeruschkamp, F.; Stephan, S.; Cui, Q.; Kotting, C.; Xia, F.; Gerwert, K. Specific substates of Ras to interact with GAPs and effectors: revealed by theoretical simulations and FTIR experiments. The journal of physical chemistry letters 2018, 9, 1312–1317.

(54) Hu, Z.; Martí, J. Atomic-level mechanisms of abnormal activation in NRAS oncogenes from two-dimensional free energy landscapes. Nanoscale 2025, 17, 4047–4057.

(55) Lu, H.; Martí, J. Predicting the conformational variability of oncogenic GTP-bound G12D mutated KRas-4B proteins at zwitterionic model cell membranes. Nanoscale 2022, 14, 3148–3158.

(56) Jurrus, E.; Engel, D.; Star, K.; Monson, K.; Brandi, J.; Felberg, L. E.; Brookes, D. H.; Wilson, L.; Chen, J.; Liles, K.; others Improvements to the APBS biomolecular solvation software suite. Protein science 2018, 27, 112–128.

(57) Dominguez, C.; Boelens, R.; Bonvin, A. M. HADDOCK: a protein-protein docking approach based on biochemical or biophysical information. Journal of the American Chemical Society 2003, 125, 1731–1737.

