## Supplementary material for "Probing the Structural Dynamics of the Unbound MAX Protein: Insights from Well-Tempered Metadynamics": reference.bib

@article{mateyak1997phenotypes,
  title={Phenotypes of c-Myc-deficient rat fibroblasts isolated by targeted homologous recombination},
  author={Mateyak, Maria K and Obaya, Alvaro J and Adachi, Susumu and Sedivy, John M},
  journal = {Cell Growth & Differentiation},
  volume = {8},
  number = {10},
  pages = {1039-1048},
  year={1997}
}
@article{luscher2001function,
  title={Function and regulation of the transcription factors of the Myc/Max/Mad network},
  author={L{\"u}scher, Bernhard},
  journal={Gene},
  volume={277},
  number={1-2},
  pages={1--14},
  year={2001},
  publisher={Elsevier}
}
@article{allevato2017sequence,
  title={Sequence-specific DNA binding by MYC/MAX to low-affinity non-E-box motifs},
  author={Allevato, Michael and Bolotin, Eugene and Grossman, Mark and Mane-Padros, Daniel and Sladek, Frances M and Martinez, Ernest},
  journal={PloS one},
  volume={12},
  number={7},
  pages={e0180147},
  year={2017},
  publisher={Public Library of Science San Francisco, CA USA}
}
@article{blackwood1991max,
  title={Max: A Helix-Loop-Helix Zipper Protein That Forms a Sequence-Specific DNA-Binding Complex with Myc},
  author={Blackwood, Elizabeth M and Eisenman, Robert N},
  journal={Science},
  volume={251},
  number={4998},
  pages={1211-1217},
  year={1991},
  publisher={American Association for the Advancement of Science}
}
@article{cascon2012max,
  title={MAX and MYC: a heritable breakup},
  author={Casc{\'o}n, Alberto and Robledo, Mercedes},
  journal={Cancer research},
  volume={72},
  number={13},
  pages={3119--3124},
  year={2012},
  publisher={American Association for Cancer Research}
}
@article{kretzner1992myc,
  title={Myc and Max proteins possess distinct transcriptional activities},
  author={Kretzner, Leo and Blackwood, Elizabeth M and Eisenman, Robert N},
  journal={Nature},
  volume={359},
  number={6394},
  pages={426--429},
  year={1992},
  publisher={Nature Publishing Group UK London}
}
@article{meyer2008reflecting,
  title={Reflecting on 25 years with MYC},
  author={Meyer, Natalie and Penn, Linda Z},
  journal={Nature Reviews Cancer},
  volume={8},
  number={12},
  pages={976--990},
  year={2008},
  publisher={Nature Publishing Group UK London}
}
@article{augert2020max,
  title={MAX functions as a tumor suppressor and rewires metabolism in small cell lung cancer},
  author={Augert, Arnaud and Mathsyaraja, Haritha and Ibrahim, Ali H and Freie, Brian and Geuenich, Michael J and Cheng, Pei-Feng and Alibeckoff, Sydney P and Wu, Nan and Hiatt, Joseph B and Basom, Ryan and others},
  journal={Cancer cell},
  volume={38},
  number={1},
  pages={97--114},
  year={2020},
  publisher={Elsevier}
}
@article{young2011premetazoan,
  title={Premetazoan ancestry of the Myc--Max network},
  author={Young, Susan L and Diolaiti, Daniel and Conacci-Sorrell, Maralice and Ruiz-Trillo, Inaki and Eisenman, Robert N and King, Nicole},
  journal={Molecular biology and evolution},
  volume={28},
  number={10},
  pages={2961--2971},
  year={2011},
  publisher={Oxford University Press}
}
@article{soucek2002omomyc,
  title={Omomyc, a potential Myc dominant negative, enhances Myc-induced apoptosis},
  author={Soucek, Laura and Jucker, Richard and Panacchia, Laura and Ricordy, Ruggero and Tato, Franco and Nasi, Sergio},
  journal={Cancer research},
  volume={62},
  number={12},
  pages={3507--3510},
  year={2002},
  publisher={American Association for Cancer Research}
}

@article{hu2025atomic,
  title={Atomic-level mechanisms of abnormal activation in NRAS oncogenes from two-dimensional free energy landscapes},
  author={Hu, Zheyao and Mart{\'\i}, Jordi},
  journal={Nanoscale},
  volume={17},
  number={7},
  pages={4047--4057},
  year={2025},
  publisher={Royal Society of Chemistry}
}

@article{lovera2015towards,
  title={Towards a molecular understanding of the link between imatinib resistance and kinase conformational dynamics},
  author={Lovera, Silvia and Morando, Maria and Pucheta-Martinez, Encarna and Martinez-Torrecuadrada, Jorge L and Saladino, Giorgio and Gervasio, Francesco L},
  journal={PLoS computational biology},
  volume={11},
  number={11},
  pages={e1004578},
  year={2015},
  publisher={Public Library of Science San Francisco, CA USA}
}

@article{barducci2008well,
  title={Well-tempered metadynamics: a smoothly converging and tunable free-energy method},
  author={Barducci, Alessandro and Bussi, Giovanni and Parrinello, Michele},
  journal={Physical review letters},
  volume={100},
  number={2},
  pages={020603},
  year={2008},
  publisher={APS}
}

@article{singh2025exploring,
  title={Exploring the Conformational Space of MPS1 Kinase Using Metadynamics},
  author={Singh, Anuradha and Lakkaniga, Naga Rajiv},
  journal={Proteins: Structure, Function, and Bioinformatics},
  volume={93},
  number={6},
  pages={1118--1127},
  year={2025},
  publisher={Wiley Online Library}
}

@article{nair2003x,
  title={X-ray structures of Myc-Max and Mad-Max recognizing DNA: molecular bases of regulation by proto-oncogenic transcription factors},
  author={Nair, Satish K and Burley, Stephen K},
  journal={Cell},
  volume={112},
  number={2},
  pages={193--205},
  year={2003},
  publisher={Elsevier}
}

@article{schutz2022disordered,
  title={The disordered MAX N-terminus modulates DNA binding of the transcription factor MYC: MAX},
  author={Sch{\"u}tz, Stefan and Bergsdorf, Christian and Goretzki, Benedikt and Lingel, Andreas and Renatus, Martin and Gossert, Alvar D and Jahnke, Wolfgang},
  journal={Journal of molecular biology},
  volume={434},
  number={22},
  pages={167833},
  year={2022},
  publisher={Elsevier}
}

@article{beaulieu2020structural,
  title={Structural and biophysical insights into the function of the intrinsically disordered Myc oncoprotein},
  author={Beaulieu, Marie-Eve and Castillo, Francisco and Soucek, Laura},
  journal={Cells},
  volume={9},
  number={4},
  pages={1038},
  year={2020},
  publisher={MDPI}
}

@article{sammak2019crystal,
  title={Crystal structures and nuclear magnetic resonance studies of the apo form of the c-MYC: MAX bHLHZip complex reveal a helical basic region in the absence of DNA},
  author={Sammak, Susan and Hamdani, Najoua and Gorrec, Fabrice and Allen, Mark D and Freund, Stefan MV and Bycroft, Mark and Zinzalla, Giovanna},
  journal={Biochemistry},
  volume={58},
  number={29},
  pages={3144--3154},
  year={2019},
  publisher={ACS Publications}
}

@article{Eyring1935,
    author = {Eyring, Henry},
    title = {The Activated Complex in Chemical Reactions},
    journal = {The Journal of Chemical Physics},
    volume = {3},
    number = {2},
    pages = {107-115},
    year = {1935},
    month = {02},
    abstract = {The calculation of absolute reaction rates is formulated in terms of quantities which are available from the potential surfaces which can be constructed at the present time. The probability of the activated state is calculated using ordinary statistical mechanics. This probability multiplied by the rate of decomposition gives the specific rate of reaction. The occurrence of quantized vibrations in the activated complex, in degrees of freedom which are unquantized in the original molecules, leads to relative reaction rates for isotopes quite different from the rates predicted using simple kinetic theory. The necessary conditions for the general statistical treatment to reduce to the usual kinetic treatment are given.},
    issn = {0021-9606},
    doi = {10.1063/1.1749604},
    url = {https://doi.org/10.1063/1.1749604},
    eprint = {https://pubs.aip.org/aip/jcp/article-pdf/3/2/107/18788362/107_1_online.pdf},
}
@article{larsen2020combining,
  title={Combining molecular dynamics simulations with small-angle X-ray and neutron scattering data to study multi-domain proteins in solution},
  author={Larsen, Andreas Haahr and Wang, Yong and Bottaro, Sandro and Grudinin, Sergei and Arleth, Lise and Lindorff-Larsen, Kresten},
  journal={PLoS computational biology},
  volume={16},
  number={4},
  pages={e1007870},
  year={2020},
  publisher={Public Library of Science San Francisco, CA USA}
}
@article{pedregosa2011scikit,
  title={Scikit-learn: Machine learning in Python},
  author={Pedregosa, Fabian and Varoquaux, Ga{\"e}l and Gramfort, Alexandre and Michel, Vincent and Thirion, Bertrand and Grisel, Olivier and Blondel, Mathieu and Prettenhofer, Peter and Weiss, Ron and Dubourg, Vincent and others},
  journal={the Journal of machine Learning research},
  volume={12},
  pages={2825--2830},
  year={2011},
  publisher={JMLR. org}
}
@misc{githubHL,
  author = {Lu, Huixia},
  title = {Project MAX},
  year = {2025},
  publisher = {GitHub},
  journal = {GitHub repository},
  howpublished = {\url{https://github.com/HuixiaLuScienceRocks/Probing_the_Structural_Dynamics_of_Unbound_MAX_Protein_Insights_from_Well_Tempered_Metadynamics}}
  }
@misc{frisch2016gaussian,
  title={Gaussian 16},
  author={Frisch, MJ ea and Trucks, GW and Schlegel, H Bernhard and Scuseria, GE and Robb, MAn and Cheeseman, JR and Scalmani, G and Barone, VPGA and Petersson, GA and Nakatsuji, HJRA and others},
  year={2016},
  publisher={Gaussian, Inc. Wallingford, CT}
}
@book{jensen2010molecular,
  title={Molecular modeling basics},
  author={Jensen, Jan H},
  year={2010},
  publisher={CRC Press}
}
@article{tomasi2005quantum,
  title={Quantum mechanical continuum solvation models},
  author={Tomasi, Jacopo and Mennucci, Benedetta and Cammi, Roberto},
  journal={Chemical reviews},
  volume={105},
  number={8},
  pages={2999--3094},
  year={2005},
  publisher={ACS Publications}
}
@article{hay1985ab,
  title={Ab initio effective core potentials for molecular calculations. Potentials for the transition metal atoms Sc to Hg},
  author={Hay, P Jeffrey and Wadt, Willard R},
  journal={The Journal of chemical physics},
  volume={82},
  number={1},
  pages={270--283},
  year={1985},
  publisher={American Institute of Physics}
}
@article{zhao2006new,
  title={A new local density functional for main-group thermochemistry, transition metal bonding, thermochemical kinetics, and noncovalent interactions},
  author={Zhao, Yan and Truhlar, Donald G},
  journal={The Journal of chemical physics},
  volume={125},
  number={19},
  year={2006},
  publisher={AIP Publishing}
}
@article{rassolov19986,
  title={6-31G* basis set for atoms K through Zn},
  author={Rassolov, Vitaly A and Pople, John A and Ratner, Mark A and Windus, Theresa L},
  journal={The Journal of chemical physics},
  volume={109},
  number={4},
  pages={1223--1229},
  year={1998},
  publisher={AIP Publishing}
}
@article{dominguez2003haddock,
  title={HADDOCK: a protein- protein docking approach based on biochemical or biophysical information},
  author={Dominguez, Cyril and Boelens, Rolf and Bonvin, Alexandre MJJ},
  journal={Journal of the American Chemical Society},
  volume={125},
  number={7},
  pages={1731--1737},
  year={2003},
  publisher={ACS Publications}
}
@article{hovsek2016metadyn,
  title={Metadyn View: Fast web-based viewer of free energy surfaces calculated by metadynamics},
  author={Ho{\v{s}}ek, Petr and Spiwok, Vojt{\v{e}}ch},
  journal={Computer Physics Communications},
  volume={198},
  pages={222--229},
  year={2016},
  publisher={Elsevier}
}

@article{brooks2009charmm,
  title={CHARMM: the biomolecular simulation program},
  author={Brooks, Bernard R and Brooks III, Charles L and Mackerell Jr, Alexander D and Nilsson, Lennart and Petrella, Robert J and Roux, Beno{\^\i}t and Won, Youngdo and Archontis, Georgios and Bartels, Christian and Boresch, Stefan and others},
  journal={Journal of computational chemistry},
  volume={30},
  number={10},
  pages={1545--1614},
  year={2009},
  publisher={Wiley Online Library}
}

@article{jo2008charmm,
  title={CHARMM-GUI: a web-based graphical user interface for CHARMM},
  author={Jo, Sunhwan and Kim, Taehoon and Iyer, Vidyashankara G and Im, Wonpil},
  journal={Journal of computational chemistry},
  volume={29},
  number={11},
  pages={1859--1865},
  year={2008},
  publisher={Wiley Online Library}
}

@article{lee2016charmm,
  title={CHARMM-GUI input generator for NAMD, GROMACS, AMBER, OpenMM, and CHARMM/OpenMM simulations using the CHARMM36 additive force field},
  author={Lee, Jumin and Cheng, Xi and Jo, Sunhwan and MacKerell, Alexander D and Klauda, Jeffery B and Im, Wonpil},
  journal={Biophysical journal},
  volume={110},
  number={3},
  pages={641a},
  year={2016},
  publisher={Elsevier}
}

@article{huang2013charmm36,
  title={CHARMM36 all-atom additive protein force field: Validation based on comparison to NMR data},
  author={Huang, Jing and MacKerell Jr, Alexander D},
  journal={Journal of computational chemistry},
  volume={34},
  number={25},
  pages={2135--2145},
  year={2013},
  publisher={Wiley Online Library}
}

@article{case2023ambertools,
  title={AmberTools},
  author={Case, David A and Aktulga, Hasan Metin and Belfon, Kellon and Cerutti, David S and Cisneros, G Andr{\'e}s and Cruzeiro, Vin{\'\i}cius Wilian D and Forouzesh, Negin and Giese, Timothy J and Gotz, Andreas W and Gohlke, Holger and others},
  journal={Journal of chemical information and modeling},
  volume={63},
  number={20},
  pages={6183--6191},
  year={2023},
  publisher={ACS Publications}
}

@article{case2005amber,
  title={The Amber biomolecular simulation programs},
  author={Case, David A and Cheatham III, Thomas E and Darden, Tom and Gohlke, Holger and Luo, Ray and Merz Jr, Kenneth M and Onufriev, Alexey and Simmerling, Carlos and Wang, Bing and Woods, Robert J},
  journal={Journal of computational chemistry},
  volume={26},
  number={16},
  pages={1668--1688},
  year={2005},
  publisher={Wiley Online Library}
}

@article{bussi2006equilibrium,
  title={Equilibrium free energies from nonequilibrium metadynamics},
  author={Bussi, Giovanni and Laio, Alessandro and Parrinello, Michele},
  journal={Physical review letters},
  volume={96},
  number={9},
  pages={090601},
  year={2006},
  publisher={APS}
}

@article{barducci2008well,
  title={Well-tempered metadynamics: a smoothly converging and tunable free-energy method},
  author={Barducci, Alessandro and Bussi, Giovanni and Parrinello, Michele},
  journal={Physical review letters},
  volume={100},
  number={2},
  pages={020603},
  year={2008},
  publisher={APS}
}

@article{bonomi2016metadynamic,
  title={Metadynamic metainference: Enhanced sampling of the metainference ensemble using metadynamics},
  author={Bonomi, Massimiliano and Camilloni, Carlo and Vendruscolo, Michele},
  journal={Scientific reports},
  volume={6},
  number={1},
  pages={31232},
  year={2016},
  publisher={Nature Publishing Group UK London}
}

@article{bonomi2009plumed,
  title={PLUMED: A portable plugin for free-energy calculations with molecular dynamics},
  author={Bonomi, Massimiliano and Branduardi, Davide and Bussi, Giovanni and Camilloni, Carlo and Provasi, Davide and Raiteri, Paolo and Donadio, Davide and Marinelli, Fabrizio and Pietrucci, Fabio and Broglia, Ricardo A and others},
  journal={Computer Physics Communications},
  volume={180},
  number={10},
  pages={1961--1972},
  year={2009},
  publisher={Elsevier}
}

@article{tribello2014plumed,
  title={PLUMED 2: New feathers for an old bird},
  author={Tribello, Gareth A and Bonomi, Massimiliano and Branduardi, Davide and Camilloni, Carlo and Bussi, Giovanni},
  journal={Computer physics communications},
  volume={185},
  number={2},
  pages={604--613},
  year={2014},
  publisher={Elsevier}
}

@article{plumed2019promoting,
  title={Promoting transparency and reproducibility in enhanced molecular simulations},
  journal={Nature methods},
  volume={16},
  number={8},
  pages={670--673},
  year={2019},
  publisher={Nature Publishing Group US New York}
}

@article{humphrey1996vmd,
  title={VMD: visual molecular dynamics},
  author={Humphrey, William and Dalke, Andrew and Schulten, Klaus},
  journal={Journal of molecular graphics},
  volume={14},
  number={1},
  pages={33--38},
  year={1996},
  publisher={Elsevier}
}

@techreport{gowers2019mdanalysis,
  title={MDAnalysis: a Python package for the rapid analysis of molecular dynamics simulations},
  author={Gowers, Richard J and Linke, Max and Barnoud, Jonathan and Reddy, Tyler John Edward and Melo, Manuel N and Seyler, Sean L and Domanski, Jan and Dotson, David L and Buchoux, S{\'e}bastien and Kenney, Ian M and others},
  year={2019},
  institution={Los Alamos National Laboratory (LANL), Los Alamos, NM (United States)}
}

@article{trapl2020analysis,
  title={Analysis of the Results of Metadynamics Simulations by metadynminer and metadynminer3d},
  author={Trapl, Dalibor and Spiwok, Vojt{\v{e}}ch},
  journal={arXiv preprint arXiv:2009.02241},
  year={2020}
}

@article{likas2003global,
  title={The global k-means clustering algorithm},
  author={Likas, Aristidis and Vlassis, Nikos and Verbeek, Jakob J},
  journal={Pattern recognition},
  volume={36},
  number={2},
  pages={451--461},
  year={2003},
  publisher={Elsevier}
}

@article{pettersen2004ucsf,
  title={UCSF Chimera—a visualization system for exploratory research and analysis},
  author={Pettersen, Eric F and Goddard, Thomas D and Huang, Conrad C and Couch, Gregory S and Greenblatt, Daniel M and Meng, Elaine C and Ferrin, Thomas E},
  journal={Journal of computational chemistry},
  volume={25},
  number={13},
  pages={1605--1612},
  year={2004},
  publisher={Wiley Online Library}
}

@article{greenacre2022principal,
  title={Principal component analysis},
  author={Greenacre, Michael and Groenen, Patrick JF and Hastie, Trevor and d’Enza, Alfonso Iodice and Markos, Angelos and Tuzhilina, Elena},
  journal={Nature Reviews Methods Primers},
  volume={2},
  number={1},
  pages={100},
  year={2022},
  publisher={Nature Publishing Group UK London}
}

@article{lu2020cellular,
  title={Cellular absorption of small molecules: free energy landscapes of melatonin binding at phospholipid membranes},
  author={Lu, Huixia and Marti, Jordi},
  journal={Scientific reports},
  volume={10},
  number={1},
  pages={9235},
  year={2020},
  publisher={Nature Publishing Group UK London}
}

@article{lu2020influence,
  title={Influence of cholesterol on the orientation of the farnesylated GTP-bound KRas-4B binding with anionic model membranes},
  author={Lu, Huixia and Mart{\'\i}, Jordi},
  journal={Membranes},
  volume={10},
  number={11},
  pages={364},
  year={2020},
  publisher={MDPI}
}

@book{janert2016gnuplot,
  title={Gnuplot in action: understanding data with graphs},
  author={Janert, Philipp K},
  year={2016},
  publisher={Simon and Schuster}
}

@article{follis2009small,
  title={Small-molecule perturbation of competing interactions between c-Myc and Max},
  author={Follis, Ariele Viacava and Hammoudeh, Dalia I and Daab, Andrew T and Metallo, Steven J},
  journal={Bioorganic \& medicinal chemistry letters},
  volume={19},
  number={3},
  pages={807--810},
  year={2009},
  publisher={Elsevier}
}

@article{hart2014inhibitor,
  title={Inhibitor of MYC identified in a Kr{\"o}hnke pyridine library},
  author={Hart, Jonathan R and Garner, Amanda L and Yu, Jing and Ito, Yoshihiro and Sun, Minghao and Ueno, Lynn and Rhee, Jin-Kyu and Baksh, Michael M and Stefan, Eduard and Hartl, Markus and others},
  journal={Proceedings of the National Academy of Sciences},
  volume={111},
  number={34},
  pages={12556--12561},
  year={2014},
  publisher={National Academy of Sciences}
}

@article{alsultan2021novel,
  title={The novel low molecular weight MYC antagonist MYCMI-6 inhibits proliferation and induces apoptosis in breast cancer cells},
  author={AlSultan, Dalal and Kavanagh, Emma and O’Grady, Shane and Eustace, Alex J and Castell, Alina and Larsson, Lars-Gunnar and Crown, John and Madden, Stephen F and Duffy, Michael J},
  journal={Investigational new drugs},
  volume={39},
  number={2},
  pages={587--594},
  year={2021},
  publisher={Springer}
}

@article{garralda2024myc,
  title={MYC targeting by OMO-103 in solid tumors: a phase 1 trial},
  author={Garralda, Elena and Beaulieu, Marie-Eve and Moreno, Victor and Casacuberta-Serra, S{\'\i}lvia and Martinez-Martin, Sandra and Foradada, Laia and Alonso, Guzman and Masso-Valles, Daniel and L{\'o}pez-Est{\'e}vez, Sergio and Jauset, Toni and others},
  journal={Nature medicine},
  volume={30},
  number={3},
  pages={762--771},
  year={2024},
  publisher={Nature Publishing Group US New York}
}

@article{henkelman2000improved,
  title={Improved tangent estimate in the nudged elastic band method for finding minimum energy paths and saddle points},
  author={Henkelman, Graeme and J{\'o}nsson, Hannes},
  journal={The Journal of chemical physics},
  volume={113},
  number={22},
  pages={9978--9985},
  year={2000},
  publisher={American Institute of Physics}
}

@article{jambeck2013exploring,
  title={Exploring the free energy landscape of solutes embedded in lipid bilayers},
  author={Jambeck, Joakim PM and Lyubartsev, Alexander P},
  journal={The Journal of Physical Chemistry Letters},
  volume={4},
  number={11},
  pages={1781--1787},
  year={2013},
  publisher={ACS Publications}
}

@article{roux1999statistical,
  title={Statistical mechanical equilibrium theory of selective ion channels},
  author={Roux, Benoit},
  journal={Biophysical journal},
  volume={77},
  number={1},
  pages={139--153},
  year={1999},
  publisher={Elsevier}
}

@article{hu2025atomic,
  title={Atomic-level mechanisms of abnormal activation in NRAS oncogenes from two-dimensional free energy landscapes},
  author={Hu, Zheyao and Mart{\'\i}, Jordi},
  journal={Nanoscale},
  volume={17},
  number={7},
  pages={4047--4057},
  year={2025},
  publisher={Royal Society of Chemistry}
}

@article{li2018specific,
  title={Specific substates of Ras to interact with GAPs and effectors: revealed by theoretical simulations and FTIR experiments},
  author={Li, Yang and Zhang, Yuwei and Gro{\ss}erüschkamp, Frederik and Stephan, Sara and Cui, Qiang and Kötting, Carsten and Xia, Fei and Gerwert, Klaus},
  journal={The journal of physical chemistry letters},
  volume={9},
  number={6},
  pages={1312--1317},
  year={2018},
  publisher={ACS Publications}
}

@article{jurrus2018improvements,
  title={Improvements to the APBS biomolecular solvation software suite},
  author={Jurrus, Elizabeth and Engel, Dave and Star, Keith and Monson, Kyle and Brandi, Juan and Felberg, Lisa E and Brookes, David H and Wilson, Leighton and Chen, Jiahui and Liles, Karina and others},
  journal={Protein science},
  volume={27},
  number={1},
  pages={112--128},
  year={2018},
  publisher={Wiley Online Library}
}


@article{lu2022predicting,
  title={Predicting the conformational variability of oncogenic GTP-bound G12D mutated KRas-4B proteins at zwitterionic model cell membranes},
  author={Lu, Huixia and Mart{\'\i}, Jordi},
  journal={Nanoscale},
  volume={14},
  number={8},
  pages={3148--3158},
  year={2022},
  publisher={Royal Society of Chemistry}
}

@article{li2018specific,
  title={Specific substates of Ras to interact with GAPs and effectors: revealed by theoretical simulations and FTIR experiments},
  author={Li, Yang and Zhang, Yuwei and Gro{\ss}erüschkamp, Frederik and Stephan, Sara and Cui, Qiang and Kötting, Carsten and Xia, Fei and Gerwert, Klaus},
  journal={The journal of physical chemistry letters},
  volume={9},
  number={6},
  pages={1312--1317},
  year={2018},
  publisher={ACS Publications}
}

@article{amati1993max,
  title={Myc and the cell cycle},
  author={Amati, Bruno and Alevizopoulos, Konstantinos and Vlach, Jamal},
  journal={Frontiers in Bioscience},
  volume={1},
  pages={d59--d67},
  year={1993}
}

@article{grandori2000max,
  title={The Myc/Max/Mad network and the transcriptional control of cell behavior},
  author={Grandori, Carla and Cowley, Shaun M and James, Lindsey P and Eisenman, Robert N},
  journal={Annual Review of Cell and Developmental Biology},
  volume={16},
  number={1},
  pages={653--699},
  year={2000},
  doi={10.1146/annurev.cellbio.16.1.653}
}

@article{luscher2001max,
  title={Function and regulation of the transcription factors of the Myc/Max/Mad network},
  author={Lüscher, Bernhard},
  journal={Gene},
  volume={277},
  number={1-2},
  pages={1--14},
  year={2001},
  doi={10.1016/S0378-1119(01)00698-1}
}

@article{sabo2014selective,
  title={Selective transcriptional regulation by Myc in cellular growth control and lymphomagenesis},
  author={Sabo, Annalisa and Kress, Thomas R and Pelizzola, Mattia and others},
  journal={Nature},
  volume={511},
  number={7510},
  pages={488--492},
  year={2014},
  doi={10.1038/nature13537}
}

@article{cole1986myc,
  title={The myc oncogene: its role in transformation and differentiation},
  author={Cole, Michael D},
  journal={Annual review of genetics},
  volume={20},
  number={1},
  pages={361--384},
  year={1986},
  publisher={Annual Reviews 4139 El Camino Way, PO Box 10139, Palo Alto, CA 94303-0139, USA}
}

@article{varmus1984molecular,
  title={The molecular genetics of cellular oncogenes},
  author={Varmus, Harold E},
  journal={Annual review of genetics},
  volume={18},
  number={1},
  pages={553--612},
  year={1984},
  publisher={Annual Reviews 4139 El Camino Way, PO Box 10139, Palo Alto, CA 94303-0139, USA}
}

@article{atchley1995myc,
  title={Myc and Max: molecular evolution of a family of proto-oncogene products and their dimerization partner.},
  author={Atchley, William R and Fitch, Walter M},
  journal={Proceedings of the National Academy of Sciences},
  volume={92},
  number={22},
  pages={10217--10221},
  year={1995},
  publisher={National Acad Sciences}
}

@article{hermeking2003myc,
  title={The MYC oncogene as a cancer drug target},
  author={Hermeking, Heiko},
  journal={Current cancer drug targets},
  volume={3},
  number={3},
  pages={163--175},
  year={2003},
  publisher={Bentham Science Publishers}
}

@article{dang2012myc,
  title={MYC on the path to cancer},
  author={Dang, Chi V},
  journal={Cell},
  volume={149},
  number={1},
  pages={22--35},
  year={2012},
  publisher={Elsevier}
}

@article{delmore2011bet,
  title={BET bromodomain inhibition as a therapeutic strategy to target c-Myc},
  author={Delmore, Jake E and Issa, Ghayas C and Lemieux, Madeleine E and Rahl, Peter B and Shi, Junwei and Jacobs, Hannah M and Kastritis, Efstathios and Gilpatrick, Timothy and Paranal, Ronald M and Qi, Jun and others},
  journal={Cell},
  volume={146},
  number={6},
  pages={904--917},
  year={2011},
  publisher={Elsevier}
}

@article{hammoudeh2009multiple,
  title={Multiple independent binding sites for small-molecule inhibitors on the oncoprotein c-Myc},
  author={Hammoudeh, Dalia I and Follis, Ariele Viacava and Prochownik, Edward V and Metallo, Steven J},
  journal={Journal of the American Chemical Society},
  volume={131},
  number={21},
  pages={7390--7401},
  year={2009},
  publisher={ACS Publications}
}

@article{kota2009therapeutic,
  title={Therapeutic microRNA delivery suppresses tumorigenesis in a murine liver cancer model},
  author={Kota, Janaiah and Chivukula, Raghu R and O'Donnell, Kathryn A and Wentzel, Erik A and Montgomery, Chrystal L and Hwang, Hun-Way and Chang, Tsung-Cheng and Vivekanandan, Perumal and Torbenson, Michael and Clark, K Reed and others},
  journal={Cell},
  volume={137},
  number={6},
  pages={1005--1017},
  year={2009},
  publisher={Elsevier}
}

@article{kiessling2006selective,
  title={Selective inhibition of c-Myc/Max dimerization and DNA binding by small molecules},
  author={Kiessling, Anke and Sperl, Bianca and Hollis, Angela and Eick, Dirk and Berg, Thorsten},
  journal={Chemistry \& biology},
  volume={13},
  number={7},
  pages={745--751},
  year={2006},
  publisher={Elsevier}
}

@article{masso2020blocking,
  title={Blocking Myc to treat cancer: reflecting on two decades of omomyc},
  author={Mass{\'o}-Vall{\'e}s, Daniel and Soucek, Laura},
  journal={Cells},
  volume={9},
  number={4},
  pages={883},
  year={2020},
  publisher={MDPI}
}

@article{beaulieu2019intrinsic,
  title={Intrinsic cell-penetrating activity propels Omomyc from proof of concept to viable anti-MYC therapy},
  author={Beaulieu, Marie-Eve and Jauset, Toni and Mass{\'o}-Vall{\'e}s, Daniel and Mart{\'\i}nez-Mart{\'\i}n, Sandra and Rahl, Peter and Maltais, Lo{\"\i}ka and Zacarias-Fluck, Mariano F and Casacuberta-Serra, S{\'\i}lvia and Serrano del Pozo, Erika and Fiore, Christopher and others},
  journal={Science translational medicine},
  volume={11},
  number={484},
  pages={eaar5012},
  year={2019},
  publisher={American Association for the Advancement of Science}
}

@article{garralda2024myc,
  title={MYC targeting by OMO-103 in solid tumors: a phase 1 trial},
  author={Garralda, Elena and Beaulieu, Marie-Eve and Moreno, V{\'\i}ctor and Casacuberta-Serra, S{\'\i}lvia and Mart{\'\i}nez-Mart{\'\i}n, Sandra and Foradada, Laia and Alonso, Guzman and Mass{\'o}-Vall{\'e}s, Daniel and L{\'o}pez-Est{\'e}vez, Sergio and Jauset, Toni and others},
  journal={Nature Medicine},
  volume={30},
  number={3},
  pages={762--771},
  year={2024},
  publisher={Nature Publishing Group US New York}
}

@article{beaulieu2020structural,
  title={Structural and biophysical insights into the function of the intrinsically disordered Myc oncoprotein},
  author={Beaulieu, Marie-Eve and Castillo, Francisco and Soucek, Laura},
  journal={Cells},
  volume={9},
  number={4},
  pages={1038},
  year={2020},
  publisher={MDPI}
}

@article{kortlever2017myc,
  title={Myc cooperates with Ras by programming inflammation and immune suppression},
  author={Kortlever, Roderik M and Sodir, Nicole M and Wilson, Catherine H and Burkhart, Deborah L and Pellegrinet, Luca and Swigart, Lamorna Brown and Littlewood, Trevor D and Evan, Gerard I},
  journal={Cell},
  volume={171},
  number={6},
  pages={1301--1315},
  year={2017},
  publisher={Elsevier}
}

@article{duffy2021myc,
  title={MYC as a target for cancer treatment},
  author={Duffy, Michael J and O'Grady, Shane and Tang, Minhong and Crown, John},
  journal={Cancer treatment reviews},
  volume={94},
  pages={102154},
  year={2021},
  publisher={Elsevier}
}

@article{lourenco2021myc,
  title={MYC protein interactors in gene transcription and cancer},
  author={Lourenco, Corey and Resetca, Diana and Redel, Cornelia and Lin, Peter and MacDonald, Alannah S and Ciaccio, Roberto and Kenney, Tristan MG and Wei, Yong and Andrews, David W and Sunnerhagen, Maria and others},
  journal={Nature Reviews Cancer},
  volume={21},
  number={9},
  pages={579--591},
  year={2021},
  publisher={Nature Publishing Group UK London}
}

@article{dhanasekaran2022myc,
  title={The MYC oncogene—the grand orchestrator of cancer growth and immune evasion},
  author={Dhanasekaran, Renumathy and Deutzmann, Anja and Mahauad-Fernandez, Wadie D and Hansen, Aida S and Gouw, Arvin M and Felsher, Dean W},
  journal={Nature reviews Clinical oncology},
  volume={19},
  number={1},
  pages={23--36},
  year={2022},
  publisher={Nature Publishing Group UK London}
}

@article{wang2021alternative,
  title={Alternative approaches to target Myc for cancer treatment},
  author={Wang, Chen and Zhang, Jiawei and Yin, Jie and Gan, Yichao and Xu, Senlin and Gu, Ying and Huang, Wendong},
  journal={Signal transduction and targeted therapy},
  volume={6},
  number={1},
  pages={117},
  year={2021},
  publisher={Nature Publishing Group UK London}
}

@article{jakobsen2024transcriptional,
  title={Transcriptional regulation by MYC: an emerging new model},
  author={Jakobsen, Simon T and Siersb{\ae}k, Rasmus},
  journal={Oncogene},
  pages={1--7},
  year={2024},
  publisher={Nature Publishing Group UK London}
}

@article{gabay2014myc,
  title={MYC activation is a hallmark of cancer initiation and maintenance},
  author={Gabay, Meital and Li, Yulin and Felsher, Dean W},
  journal={Cold Spring Harbor perspectives in medicine},
  volume={4},
  number={6},
  pages={a014241},
  year={2014},
  publisher={Cold Spring Harbor Laboratory Press}
}
@article{Larsen2012,
author = {Lindorff-Larsen, Kresten and Trbovic, Nikola and Maragakis, Paul and Piana, Stefano and Shaw, David E.},
title = {Structure and Dynamics of an Unfolded Protein Examined by Molecular Dynamics Simulation},
journal = {Journal of the American Chemical Society},
volume = {134},
number = {8},
pages = {3787-3791},
year = {2012},
doi = {10.1021/ja209931w},
}
