## Supplementary material for "Probing the Structural Dynamics of the Unbound MAX Protein: Insights from Well-Tempered Metadynamics": SI.pdf

### Supporting Information Available

#### Further details on PCA analysis

Here we provide further details on the principal component analysis reported in the main paper. Fig. S1 shows the variance and cumulative variance of the data explained by up to 10 PC variables, justifying that the first few components capture the essential molecular motion. Fig. S2 presents an alternative clustering of the PCA results using 8 clusters instead of the 7 clusters employed in the main text. As shown in that figure, increasing the number of clusters does not add a significantly different new structure and the additional cluster has lower density (fraction of configurations in that cluster) than the other clusters. The overall clustering pattern closely mirrors the 7-cluster analysis presented in the main text, with the additional cluster reflecting a finer subdivision within the primary group. These results support the conclusion that 7 clusters are sufficient to capture the dominant structural features of the dataset.

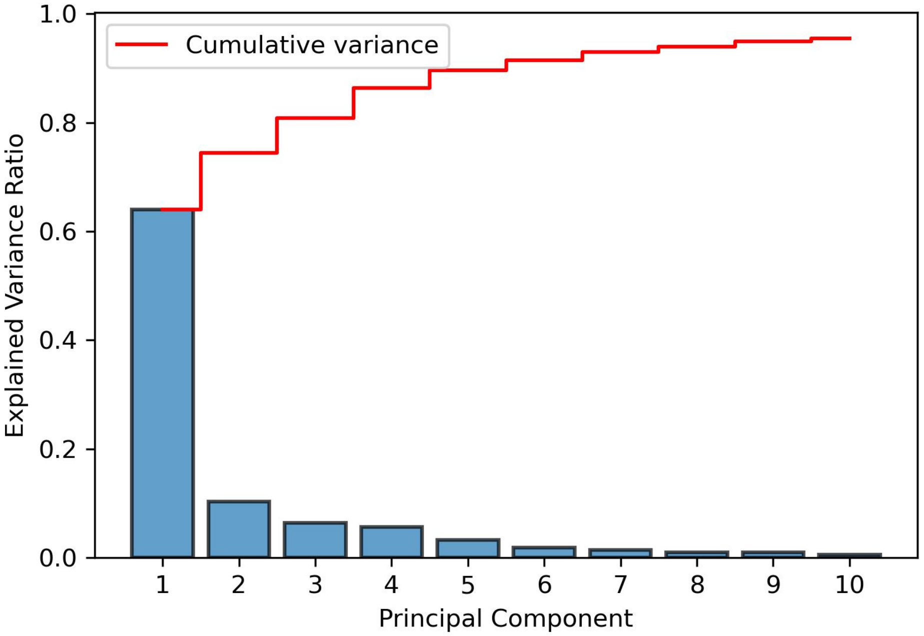


Figure S1: Percentage of explained variance for the first 10 principal components (blue bars) and cumulative variance (red).

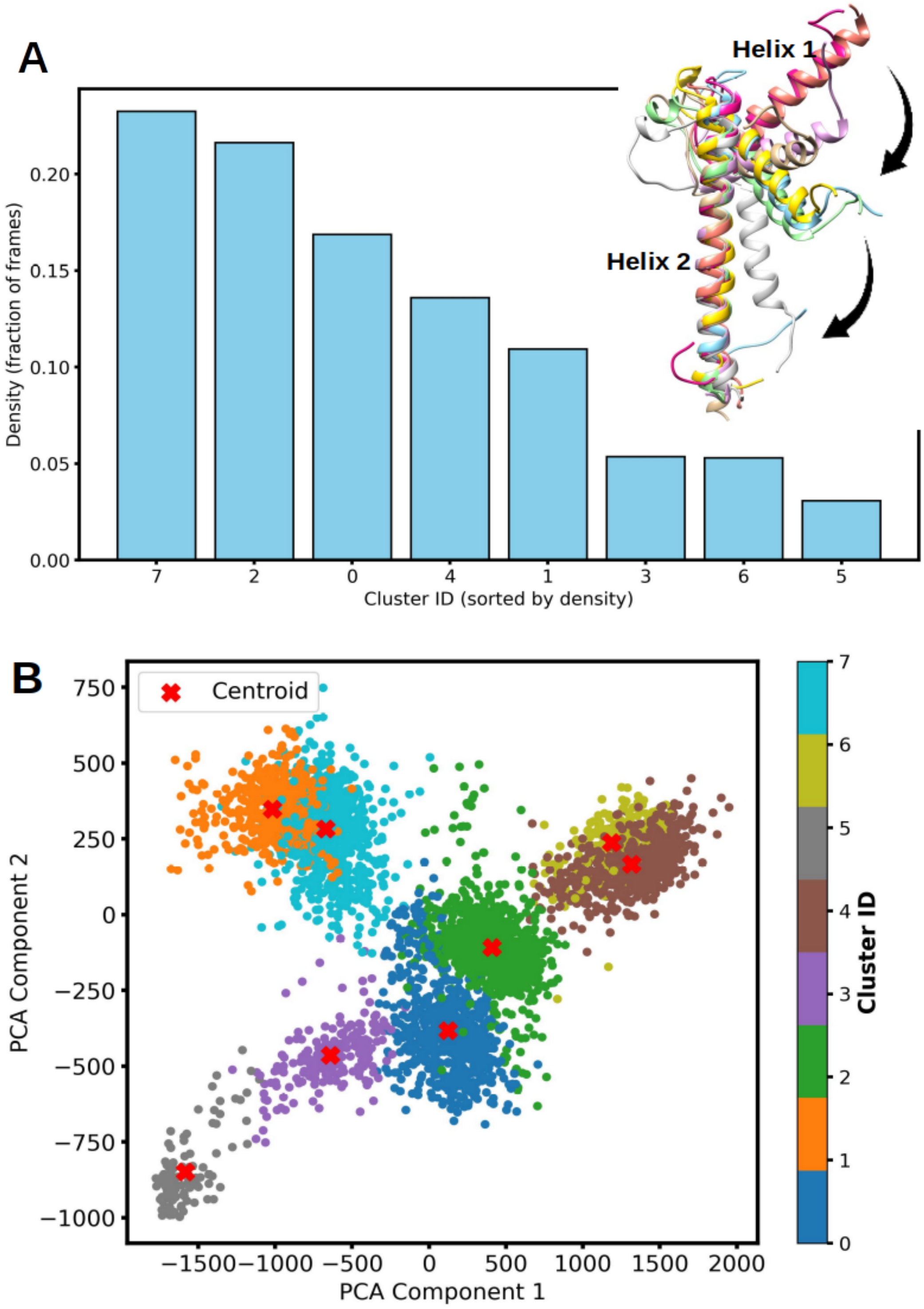


Figure S2: PCA-based clustering results using 8 clusters. (A) Density profiles of the identified clusters. (B) 2D plot of the clustering of the PCA data in 8 clusters, with representative structures marked by red crosses. The protein structures shown in the figure correspond to the representative structures for each cluster (superimposed), shown using the same color employed in the clustering plot.

### Convergence of WT-MetaD simulations

The Gaussian kernels employed during the simulation are shown in Fig. S3. This figure shows that as the simulation progresses, the bias potential gradually fills the underlying free energy surface, enabling the system to explore the full phase space. A general downward trend in the height of the employed bias is evident, becoming noticeably flatter around 1600 ns, with only minor fluctuations. In Fig. S4 we show the values of the two collective variables explored during the simulation. The figure shows that the selected two-dimensional collective variable space has been comprehensively sampled along the simulation time. In order to monitor possible folding or unfolding events during the metadynamics simulation, the proportion of secondary helices in the MAX secondary structure throughout the entire WT-metaD simulation time was determined using the STRIDE plugin within VMD, see Fig. S5. Between 500-3500 ns, we observe the fluctuations between reversible folding (helicity increasing up to $\sim 45$ %) and unfolding (helicity decreasing to $\sim 22$ %), following an initial drop in helicity from the initial crystal structure. The previous results suggest that the WT-MetaD simulation has correctly sampled the targeted configuration space. We now assess the convergence of the free energy calculations. To this end, we employ two widely used and reliable criteria: (1) comparing free energy estimates as a function of each CV at different simulation time, and (2) performing block analysis to estimate the statistical error in the free energy profiles. From the results of Fig. S6 (a) and (b), we can see that the free energy profile does not change significantly as a function of time, which indicates that the system most likely converged for the two CVs considered in the present study. In all cases, very well-converged profiles can be observed around 3350 ns, regardless of the type of CV.

The average error in the free energy profiles is computed and presented in Fig. S7. As expected, the errors increase with the block length until they reach a plateau in all cases. The average errors tend to stabilize around 0.72 kJ/mol for *d1* and 1.06 kJ/mol for *d2*, indicating consistent behavior over extended simulation times. Both criteria indicate that the well-tempered metadynamics simulations performed in this study have converged.

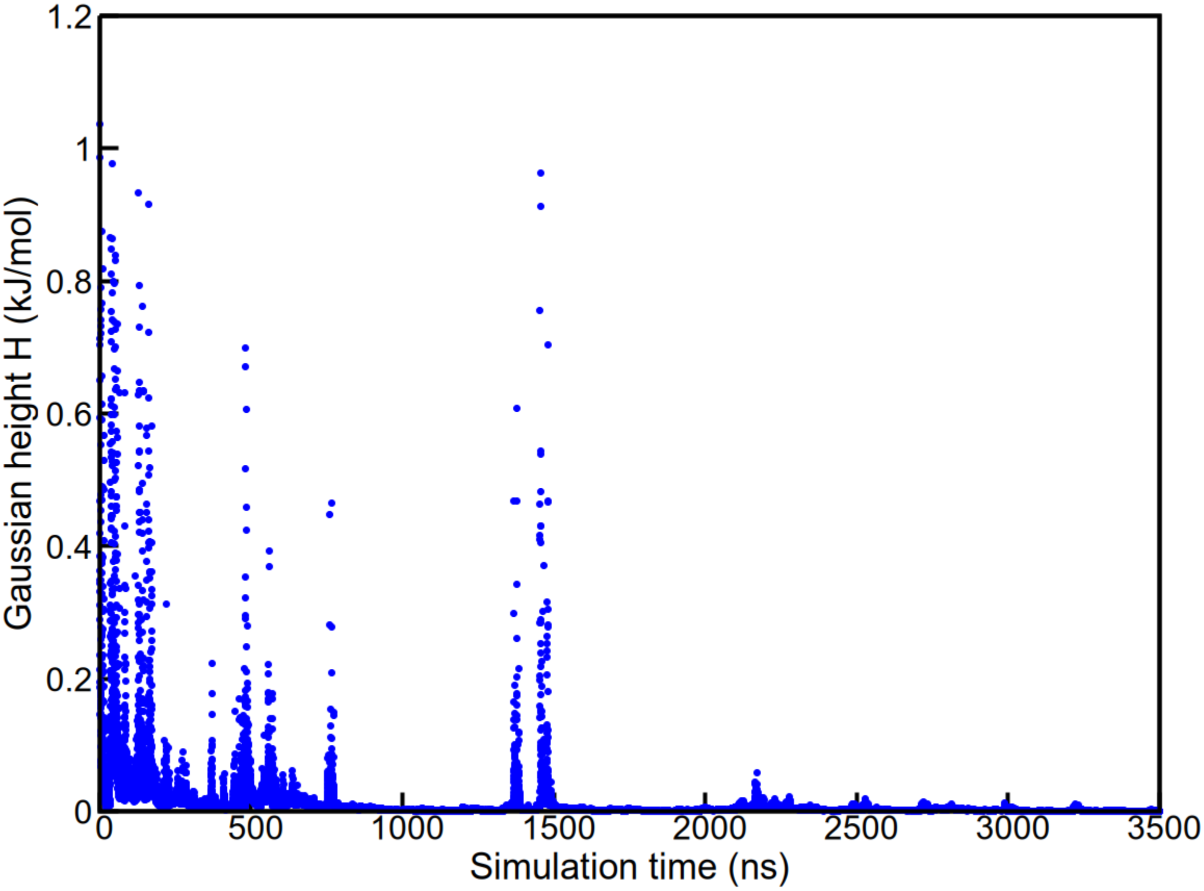


Figure S3: Time-evolution of the Gaussian height H during the overall simulation time of WT-MetaD.

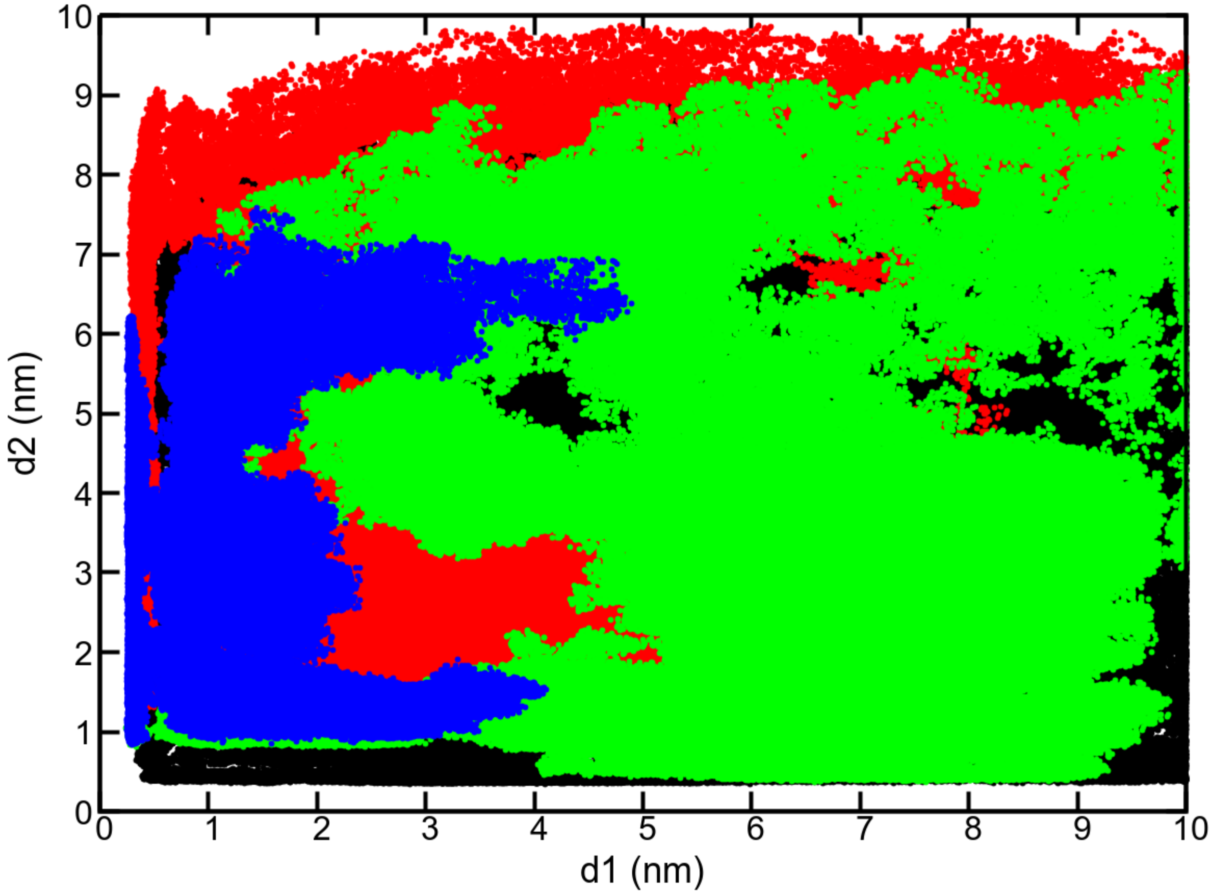


Figure S4: 2D distribution of the two collective variables sampled throughout the entire WT-MetaD simulation. Regions explored within 0-1000 $\mu$s (black), 1000-2000 $\mu$s (red), 2000-3000 $\mu$s (green), and 3000-3500 $\mu$s (blue).

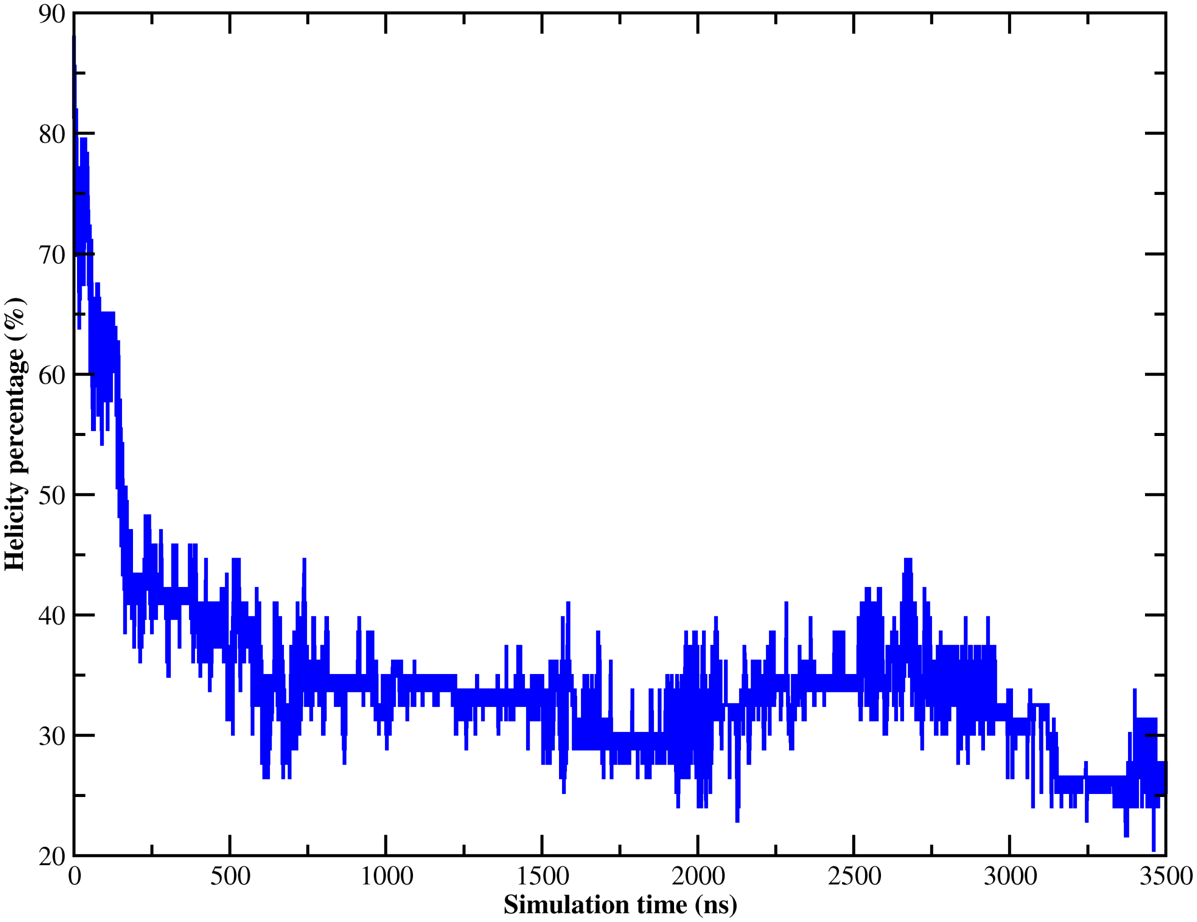


Figure S5: Helicity percentage in the secondary structure of MAX along the entire WT-MetaD simulation.

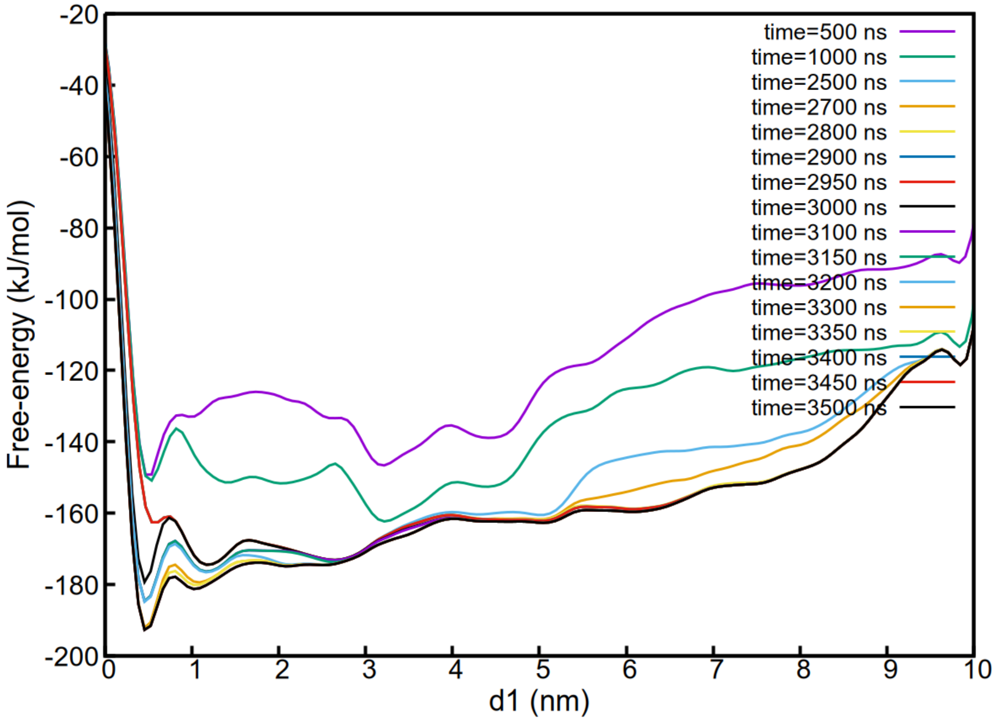


(a) Estimates of the free energy as a function of *d1*.

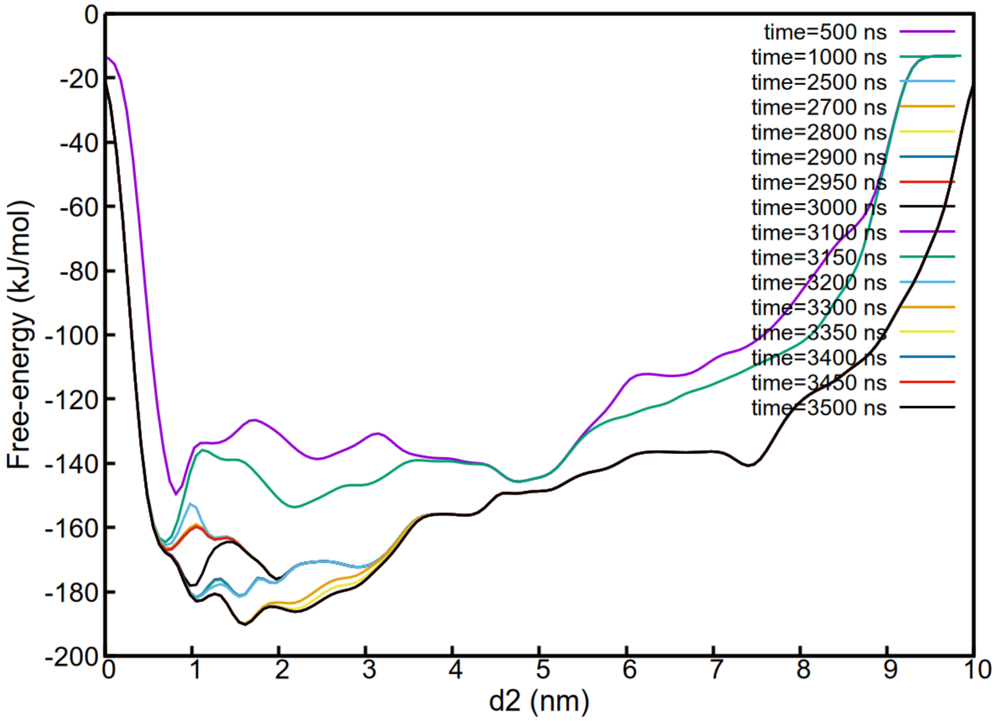


(b) Estimates of the free energy as a function of *d2*.

Figure S6: Convergence of one-dimensional free energies at different simulation times.

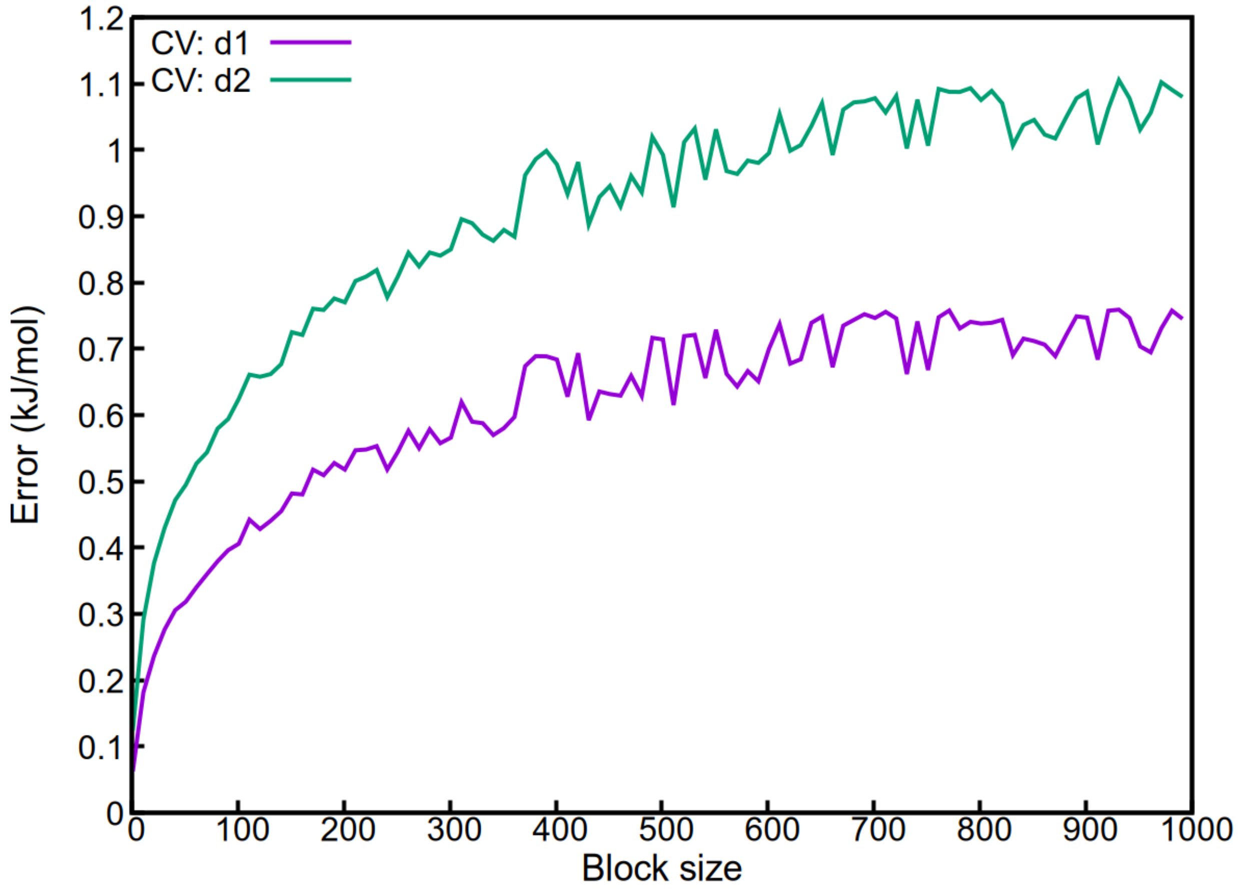


Figure S7: Block analysis profile using *d1* and *d2*.

### Free energy profiles along the Minimum Free Energy Paths

In the main text, we report the minimum free energy paths connecting different free energy basins as identified by the Nudged Elastic Band method[1] (Fig. 4). Although the free energy values can be identified from that figure, in order to facilitate the discussion, we show here the free energy profiles explicitly evaluated over these paths (Fig. S8). We also indicate the transition states found along these paths.

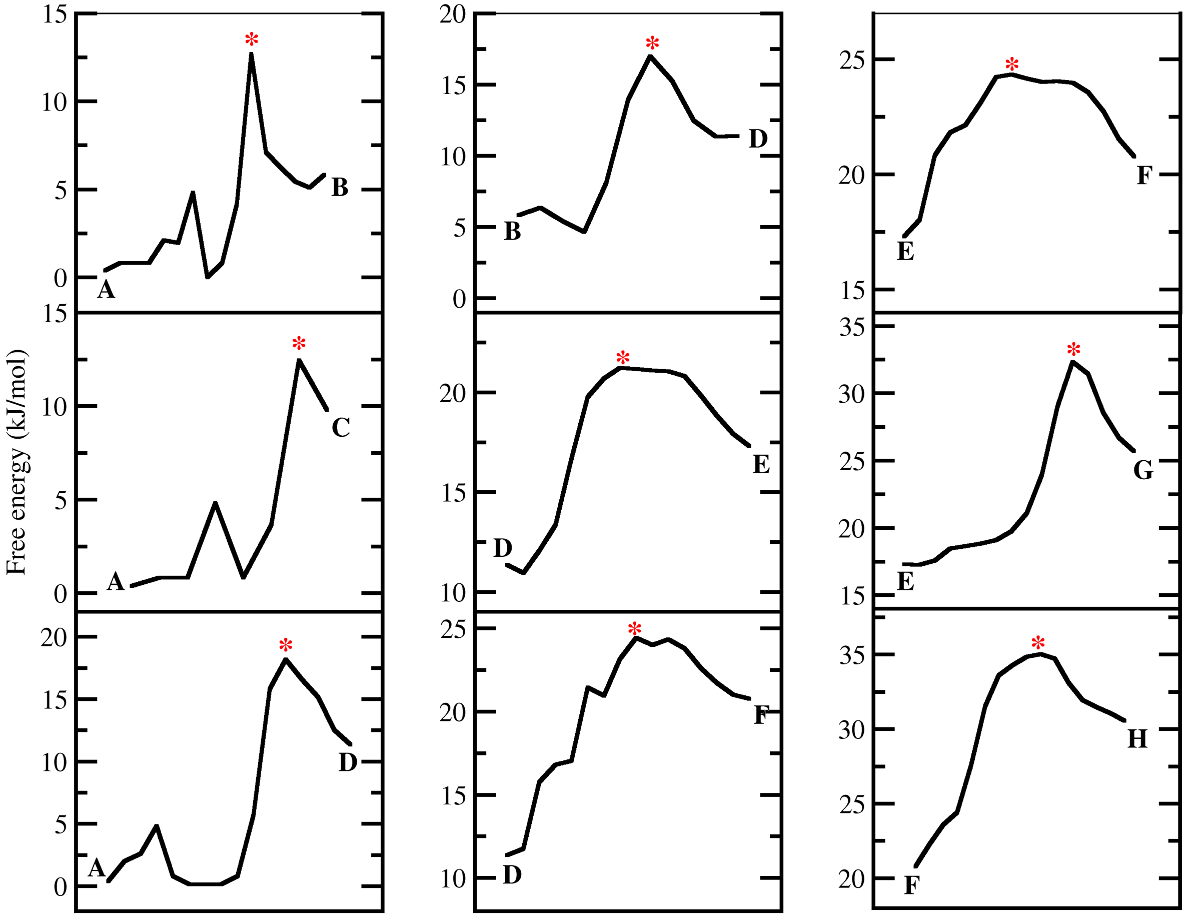


Figure S8: Free Energy profiles evaluated over the minimum free energy paths shown in Fig. 4 of the main text. The transition states are indicated with a red star.

### Structural optimization of the inhibitor 10058-F4

Before conducting a docking analysis of the inhibitor 10058-F4 to possible drug pockets proposed in our work, we first performed density functional theory (DFT) calculations to determine the optimized geometry of this molecule and the most energetically favorable form in water solvent using Gaussian v16 software.[2] We used the CHARMM36m force field–predicted structure as the initial model for the ligand 10058-F4. For the DFT geometry optimization, a hybrid functional with a standard basis set was employed: B3LYP/6-31G*.[3,4] We used a polarizable continuum model (PCM) to represent the water using the integral equation formalism variant (IEFPCM).[5] The energy difference obtained after a geometry optimization for the relaxed 10058-F4 is -15.54 kcal/mol compared to the initial structure predicted by CHARMM36m force field. Their corresponding structures are reported in Fig. S9.

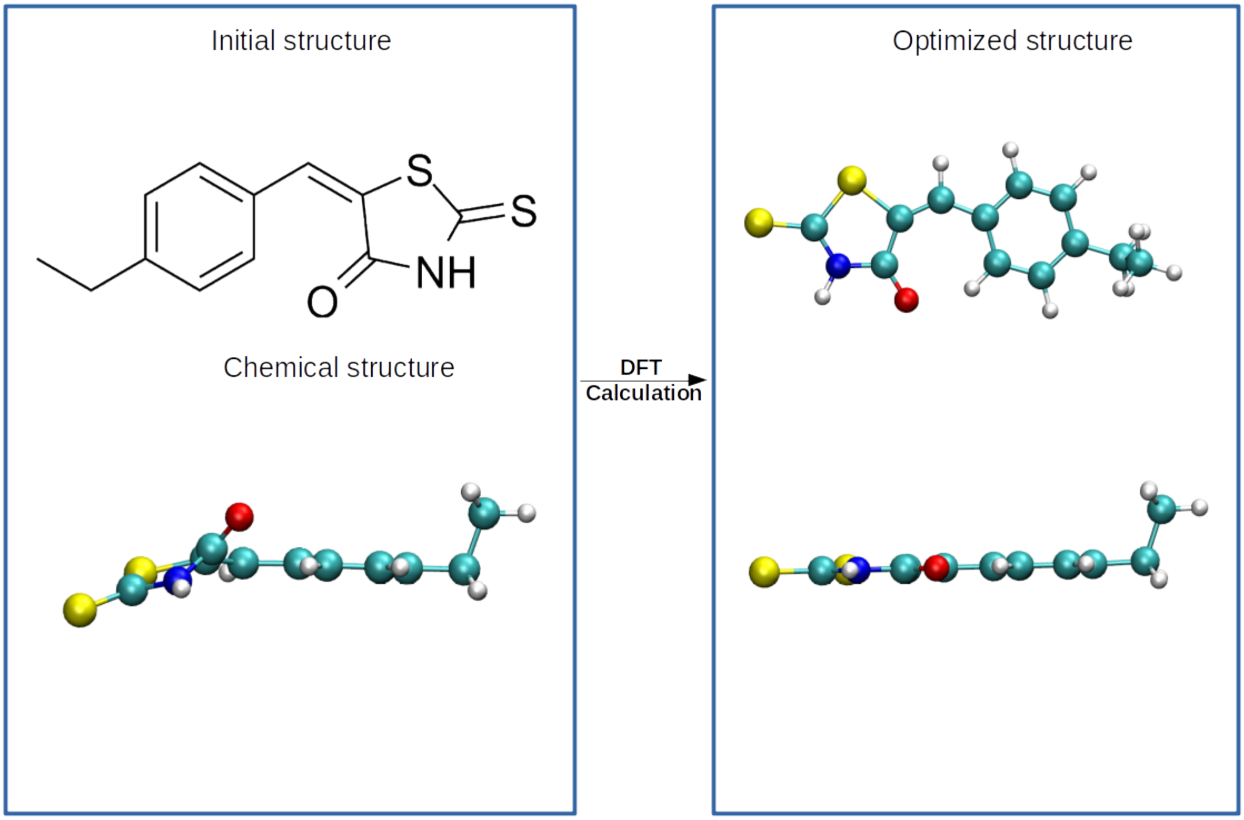


Figure S9: Initial structure and optimized structure of the inhibitor 10058-F4.

#### Inhibitor-MAX docking simulations

HADDOCK (High Ambiguity Driven biomolecular DOCKing)[6] is a docking tool that allows for the flexible docking of different states of unbound MAX to inhibitors. All residues of MAX were selected as the active binding sites for this inhibitor. Non-polar hydrogen atoms were deleted except those bonded to a polar atom (N, O atoms) before the docking procedure. A fraction (2 %) of the ambiguous restraints was randomly excluded to account for conformational flexibility. Initial docking included 1000 structures for rigid body docking, followed by 5 trials of rigid body minimization. A final stage of short molecular dynamics in explicit water solvent was adopted. Then, 200 structures were selected for the semi-flexible refinement, and a final refinement was conducted. The fraction of Common Contacts was adopted for clustering and the minimum cluster size of 4 was selected during the docking process. Cross-docking with randomized starting orientations was performed to ensure a diverse sampling of conformational space.

### References


(1) Henkelman, G.; Jónsson, H. Improved tangent estimate in the nudged elastic band method for finding minimum energy paths and saddle points. *The Journal of chemical physics* **2000**, *113*, 9978–9985.

(2) Frisch, M. e.; Trucks, G.; Schlegel, H. B.; Scuseria, G.; Robb, M.; Cheeseman, J.; Scalmani, G.; Barone, V.; Petersson, G.; Nakatsuji, H.; others Gaussian 16. 2016.

(3) Jensen, J. H. *Molecular modeling basics*; CRC Press, 2010.

(4) Rassolov, V. A.; Pople, J. A.; Ratner, M. A.; Windus, T. L. 6-31G* basis set for atoms K through Zn. *The Journal of chemical physics* **1998**, *109*, 1223–1229.

(5) Tomasi, J.; Mennucci, B.; Cammi, R. Quantum mechanical continuum solvation models. *Chemical reviews* **2005**, *105*, 2999–3094.

(6) Dominguez, C.; Boelens, R.; Bonvin, A. M. HADDOCK: a protein- protein docking approach based on biochemical or biophysical information. *Journal of the American Chemical Society* **2003**, *125*, 1731–1737.
