## Supplementary material for "Probing the Structural Dynamics of the Unbound MAX Protein: Insights from Well-Tempered Metadynamics": supporting_info.tex

%%%%%%%%%%%%%%%%%%%%%%%%%%%%%%%%%%%%%%%%%%%%%%%%%%%%%%%%%%%%%%%%%%%%%
%% This is a (brief) model paper using the achemso class
%% The document class accepts keyval options, which should include
%% the target journal and optionally the manuscript type. 
%%%%%%%%%%%%%%%%%%%%%%%%%%%%%%%%%%%%%%%%%%%%%%%%%%%%%%%%%%%%%%%%%%%%%
\documentclass[journal=jacsat,manuscript=article]{achemso}

%%%%%%%%%%%%%%%%%%%%%%%%%%%%%%%%%%%%%%%%%%%%%%%%%%%%%%%%%%%%%%%%%%%%%
%% Place any additional packages needed here.  Only include packages
%% which are essential, to avoid problems later. Do NOT use any
%% packages which require e-TeX (for example etoolbox): the e-TeX
%% extensions are not currently available on the ACS conversion
%% servers.
%%%%%%%%%%%%%%%%%%%%%%%%%%%%%%%%%%%%%%%%%%%%%%%%%%%%%%%%%%%%%%%%%%%%%
\usepackage[version=3]{mhchem} % Formula subscripts using \ce{}
\usepackage{multirow}
\usepackage[table]{xcolor}

\usepackage{placeins}
%\usepackage{hyperref}
%%%%%%%%%%%%%%%%%%%%%%%%%%%%%%%%%%%%%%%%%%%%%%%%%%%%%%%%%%%%%%%%%%%%%
%% If issues arise when submitting your manuscript, you may want to
%% un-comment the next line.  This provides information on the
%% version of every file you have used.
%%%%%%%%%%%%%%%%%%%%%%%%%%%%%%%%%%%%%%%%%%%%%%%%%%%%%%%%%%%%%%%%%%%%%
%%\listfiles
%Image-related packages
\usepackage{graphicx}
\usepackage{subcaption}
\usepackage[export]{adjustbox}
\usepackage{wrapfig}
%%%%%%%%%%%%%%%%%%%%%%%%%%%%%%%%%%%%%%%%%%%%%%%%%%%%%%%%%%%%%%%%%%%%%
%% Place any additional macros here.  Please use \newcommand* where
%% possible, and avoid layout-changing macros (which are not used
%% when typesetting).
%%%%%%%%%%%%%%%%%%%%%%%%%%%%%%%%%%%%%%%%%%%%%%%%%%%%%%%%%%%%%%%%%%%%%
\newcommand*\mycommand[1]{\texttt{\emph{#1}}}
\renewcommand{\thefigure}{S\arabic{figure}}

\usepackage{caption}

\renewcommand{\thetable}{S\arabic{table}}
\renewcommand{\thepage}{S\arabic{page}}

%%%%%%%%%%%%%%%%%%%%%%%%%%%%%%%%%%%%%%%%%%%%%%%%%%%%%%%%%%%%%%%%%%%%%
%% Meta-data block
%% ---------------
%% Each author should be given as a separate \author command.
%%
%% Corresponding authors should have an e-mail given after the author
%% name as an \email command. Phone and fax numbers can be given
%% using \phone and \fax, respectively; this information is optional.
%%
%% The affiliation of authors is given after the authors; each
%% \affiliation command applies to all preceding authors not already
%% assigned an affiliation.
%%
%% The affiliation takes an option argument for the short name.  This
%% will typically be something like "University of Somewhere".
%%
%% The \altaffiliation macro should be used for new address, etc.
%% On the other hand, \alsoaffiliation is used on a per author basis
%% when authors are associated with multiple institutions.
%%%%%%%%%%%%%%%%%%%%%%%%%%%%%%%%%%%%%%%%%%%%%%%%%%%%%%%%%%%%%%%%%%%%%
\author{Huixia Lu}
\affiliation[Institut de Ci$\'e$ncia de Materials de Barcelona (ICMAB-CSIC)]
{Institut de Ciència de Materials de Barcelona (ICMAB-CSIC), \textcolor{black}{Campus de la UAB, E-08193 Bellaterra, Spain}}        
\author{Jordi Marti}
\affiliation[Universitat Polit$\'e$cnica de Catalunya-Barcelona Tech (UPC)]
{Department of Physics, Universitat Politècnica de Catalunya-Barcelona Tech (UPC), Barcelona, Spain}
\author{Jordi Faraudo}
\affiliation[Institut de Ci$\'e$ncia de Materials de Barcelona (ICMAB-CSIC)]
{Institut de Ciència de Materials de Barcelona (ICMAB-CSIC), \textcolor{black}{Campus de la UAB, E-08193 Bellaterra, Spain}} 
%%%%%%%%%%%%%%%%%%%%%%%%%%%%%%%%%%%%%%%%%%%%%%%%%%%%%%%%%%%%%%%%%%%%%
%% The document title should be given as usual. Some journals require
%% a running title from the author: this should be supplied as an
%% optional argument to \title.
%%%%%%%%%%%%%%%%%%%%%%%%%%%%%%%%%%%%%%%%%%%%%%%%%%%%%%%%%%%%%%%%%%%%%
\title{Supporting information: Probing the Structural Dynamics of the Unbound MAX Protein: Insights from Well-Tempered Metadynamics}


\begin{document}

%%%%%%%%%%%%%%%%%%%%%%%%%%%%%%%%%%%%%%%%%%%%%%%%%%%%%%%%%%%%%%%%%%%%%
%% The "tocentry" environment can be used to create an entry for the
%% graphical table of contents. It is given here as some journals
%% require that it is printed as part of the abstract page. It will
%% be automatically moved as appropriate.
%%%%%%%%%%%%%%%%%%%%%%%%%%%%%%%%%%%%%%%%%%%%%%%%%%%%%%%%%%%%%%%%%%%%%
%\begin{tocentry}

%\begin{figure}[!h]
%\centering
%\includegraphics[width=0.85\textwidth]{toc.pdf}
%\caption{ } 
%\label{toc}
%\end{figure}

%\end{tocentry}

%%%%%%%%%%%%%%%%%%%%%%%%%%%%%%%%%%%%%%%%%%%%%%%%%%%%%%%%%%%%%%%%%%%%%
%% The abstract environment will automatically gobble the contents
%% if an abstract is not used by the target journal.
%%%%%%%%%%%%%%%%%%%%%%%%%%%%%%%%%%%%%%%%%%%%%%%%%%%%%%%%%%%%%%%%%%%%%
\begin{suppinfo}

\subsection*{Further details on PCA analysis}

\textcolor{black}{Here we provide further details on the principal component analysis reported in the main paper. Fig. \ref{fig1-si} shows the variance and cumulative variance of the data explained by up to 10 PC variables, justifying that the first few components capture the essential molecular motion.
Fig. \ref{fig2-si} presents an alternative clustering of the PCA results using 8 clusters instead of the 7 clusters employed in the main text. As shown in that figure, increasing the number of clusters does not add a significantly different new structure and the additional cluster has lower density (fraction of configurations in that cluster) than the other clusters.}
The overall clustering pattern closely mirrors the \textcolor{black}{7}-cluster analysis presented in the main text, with the additional cluster reflecting a finer subdivision within the primary group. These results support the conclusion that \textcolor{black}{7} clusters are sufficient to capture the dominant structural features of the dataset.

\begin{figure}[!h]
\centering
\includegraphics[width=0.85\textwidth]{SI-fig1.jpg}
\caption{Percentage of explained variance for the first 10 principal components (blue bars) and  cumulative variance (red).} 
\label{fig1-si}
\end{figure}

\begin{figure}[!h]
\centering
\includegraphics[width=0.85\textwidth]{SI-fig2.png}
\caption{PCA-based clustering results using 8 clusters. (A) Density profiles of the identified clusters.
(B) 2D plot of the clustering of the PCA data in 8 clusters, with representative structures marked by red crosses. The protein structures shown in the figure correspond to the representative structures for each cluster (superimposed), shown using the same color employed in the clustering plot.} 
\label{fig2-si}
\end{figure}

\FloatBarrier

\subsection*{Convergence of WT-MetaD simulations}
The Gaussian kernels employed during the simulation are shown in Fig. \ref{fig3-si}. This figure shows that as the simulation progresses, the bias potential gradually fills the underlying free energy surface, enabling the system to explore the full phase space. A general downward trend in the height of the employed bias is evident, becoming noticeably flatter around 1600 ns, with only minor fluctuations. In Fig. \ref{fig4-si} we show the values of the two collective variables explored during the simulation. 
The figure shows that the selected two-dimensional collective variable space has been comprehensively sampled along the simulation time.
In order to monitor possible folding or unfolding events during the metadynamics simulation, the proportion of secondary helices in the MAX secondary structure throughout the entire WT-metaD simulation time was determined using the STRIDE plugin within VMD, see Fig.~\ref{helicity}. Between 500-3500 ns, we observe the fluctuations between reversible folding (helicity increasing up to $\sim$ 45 $\%$) and unfolding (helicity decreasing to $\sim$ 22 $\%$), following an initial drop in helicity from the initial crystal structure. 
The previous results suggest that the WT-MetaD simulation has correctly sampled the targeted configuration space. We now assess the convergence of the free energy calculations. To this end, we employ two widely used and reliable criteria: (1) comparing free energy estimates as a function of each CV at different simulation time, and (2) performing block analysis to estimate the statistical error in the free energy profiles. From the results of Fig. \ref{fig5-si} (a) and (b), we can see that the free energy profile does not change significantly as a function of time, which indicates that the system most likely converged for the two CVs considered in the present study. In all cases, very well-converged profiles can be observed around 3350 ns, regardless of the type of CV.
\begin{figure}[hbt!]
\includegraphics[width=1.0\linewidth]{SI-fig3.png}
\caption{Time-evolution of the Gaussian height H during the overall simulation time of WT-MetaD.}
\label{fig3-si}
\end{figure}

\begin{figure}[hbt!]
\includegraphics[width=1.\linewidth]{SI-fig4.png}
\caption{2D distribution of the two collective variables sampled throughout the entire WT-MetaD simulation.
\textcolor{black}{Regions explored within 0-1000 $\mu$s (black), 1000-2000 $\mu$s (red), 2000-3000 $\mu$s (green), and 3000-3500 $\mu$s (blue).}}
\label{fig4-si}
\end{figure}

\begin{figure}[hbt!]
\includegraphics[width=1.\linewidth]{SI-helicity.png}
\caption{\textcolor{black}{Helicity percentage in the secondary structure of MAX along the entire WT-MetaD simulation.}}
\label{helicity}
\end{figure}

\begin{figure}[h]
\begin{subfigure}{0.9\textwidth}
\centering
\includegraphics[width=0.9\linewidth, height=10cm]{SI-fig5-a.png} 
\caption{Estimates of the free energy as a function of \textit{d1}.}
\label{fig:subim1}
\end{subfigure}
\begin{subfigure}{0.9\textwidth}
\centering
\includegraphics[width=0.9\linewidth, height=10cm]{SI-fig5-b.png}
\caption{Estimates of the free energy as a function of \textit{d2}.}
\label{fig:subim2}
\end{subfigure}
\caption{\textcolor{black}{Convergence of one-dimensional free energies} at different simulation times.}
\label{fig5-si}
\end{figure}

The average error in the free energy profiles is computed and presented in Fig.~\ref{fig6-si}. As expected, the errors increase with the block length until they reach a plateau in all cases. The average errors tend to stabilize around 0.72 kJ/mol for \textit{d1} and 1.06 kJ/mol for \textit{d2}, indicating consistent behavior over extended simulation times. Both criteria indicate that the well-tempered metadynamics simulations performed in this study have converged.

\begin{figure}[hbt!]
\includegraphics[width=1.\linewidth]{SI-fig6.png}
\caption{Block analysis profile using \textit{d1} and \textit{d2}.}
\label{fig6-si}
\end{figure}

\FloatBarrier

\subsection*{\textcolor{black}{Free energy profiles along the Minimum Free Energy Paths}}

\textcolor{black}{
In the main text, we report the minimum free energy paths connecting different free energy basins as identified by the Nudged Elastic Band method\cite{henkelman2000improved} (Fig. 4). 
Although the free energy values can be identified from that figure, in order to facilitate the discussion, we show here the free energy profiles explicitly evaluated over these paths (Fig.~\ref{fig7-si}). 
We also indicate the transition states found along these paths. }

\begin{figure}[hbt!]
\includegraphics[width=1.\linewidth]{SI-fig7.png}
\caption{Free Energy profiles evaluated over the minimum free energy paths shown in Fig. 4 of the main text. The transition states are indicated with a red star.}
\label{fig7-si}
\end{figure}

\FloatBarrier

\subsection*{\textcolor{black}{Structural optimization of the inhibitor 10058-F4}}
\textcolor{black}{Before conducting a docking analysis of the inhibitor 10058-F4 to possible drug pockets proposed in our work, we first performed density functional theory (DFT) calculations to determine the optimized geometry of this molecule and the most energetically favorable form in water solvent using Gaussian v16 software\cite{frisch2016gaussian}. We used the CHARMM36m force field–predicted structure as the initial model for the ligand 10058-F4.
For the DFT geometry optimization, a hybrid functional with a standard basis set was employed: B3LYP/6-31G*\cite{jensen2010molecular,rassolov19986}.
We used a polarizable continuum model (PCM) to represent the water using the integral equation formalism variant (IEFPCM)\cite{tomasi2005quantum}.
The energy difference obtained after a geometry optimization for the relaxed 10058-F4 is -15.54 kcal/mol compared to the initial structure predicted by CHARMM36m force field. Their corresponding structures are reported in Fig.~\ref{fig8-si}.}
\begin{figure}[hbt!]
\includegraphics[width=1.\linewidth]{SI-fig8.png}
\caption{Initial structure and optimized structure of the inhibitor 10058-F4.}
\label{fig8-si}
\end{figure}

\FloatBarrier
\subsection*{Inhibitor-MAX docking simulations}
HADDOCK (High Ambiguity Driven biomolecular DOCKing)\cite{dominguez2003haddock} is a docking tool that allows for the flexible docking of different states of unbound MAX to inhibitors.
All residues of MAX were selected as the active binding sites for this inhibitor. Non-polar hydrogen atoms were deleted except those bonded to a polar atom (N, O atoms) before the docking procedure. A fraction (2~$\%$) of the ambiguous restraints was randomly excluded to account for conformational flexibility. Initial docking included 1000 structures for rigid body docking, followed by 5 trials of rigid body minimization. A final stage of short molecular dynamics in explicit water solvent was adopted. Then, 200 structures were selected for the semi-flexible refinement, and a final refinement was conducted. The fraction of Common Contacts was adopted for clustering and the minimum cluster size of 4 was selected during the docking process.
Cross-docking with randomized starting orientations was performed to ensure a diverse sampling of conformational space.

\FloatBarrier
\end{suppinfo}
%%%%%%%%%%%%%%%%%%%%%%%%%%%%%%%%%%%%%%%%%%%%%%%%%%%%%%%%%%%%%%%%%%%%%
%% The appropriate \bibliography command should be placed here.
%% Notice that the class file automatically sets \bibliographystyle
%% and also names the section correctly.
%%%%%%%%%%%%%%%%%%%%%%%%%%%%%%%%%%%%%%%%%%%%%%%%%%%%%%%%%%%%%%%%%%%%%

\bibliography{reference}

\end{document}
