## Supplementary figures and images for "Probing the Structural Dynamics of the Unbound MAX Protein: Insights from Well-Tempered Metadynamics"

### SI-fig1.jpg

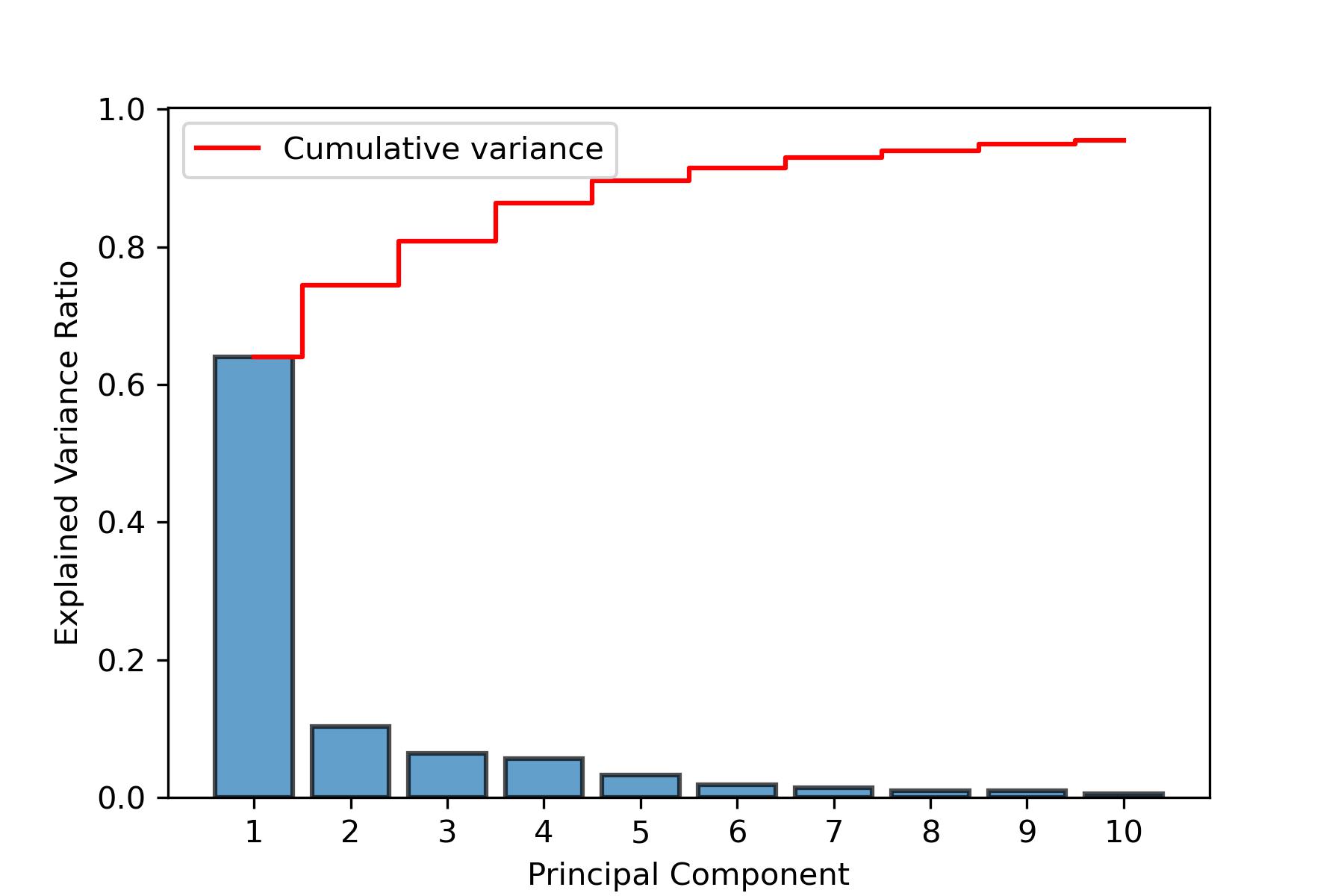

### SI-fig2.png

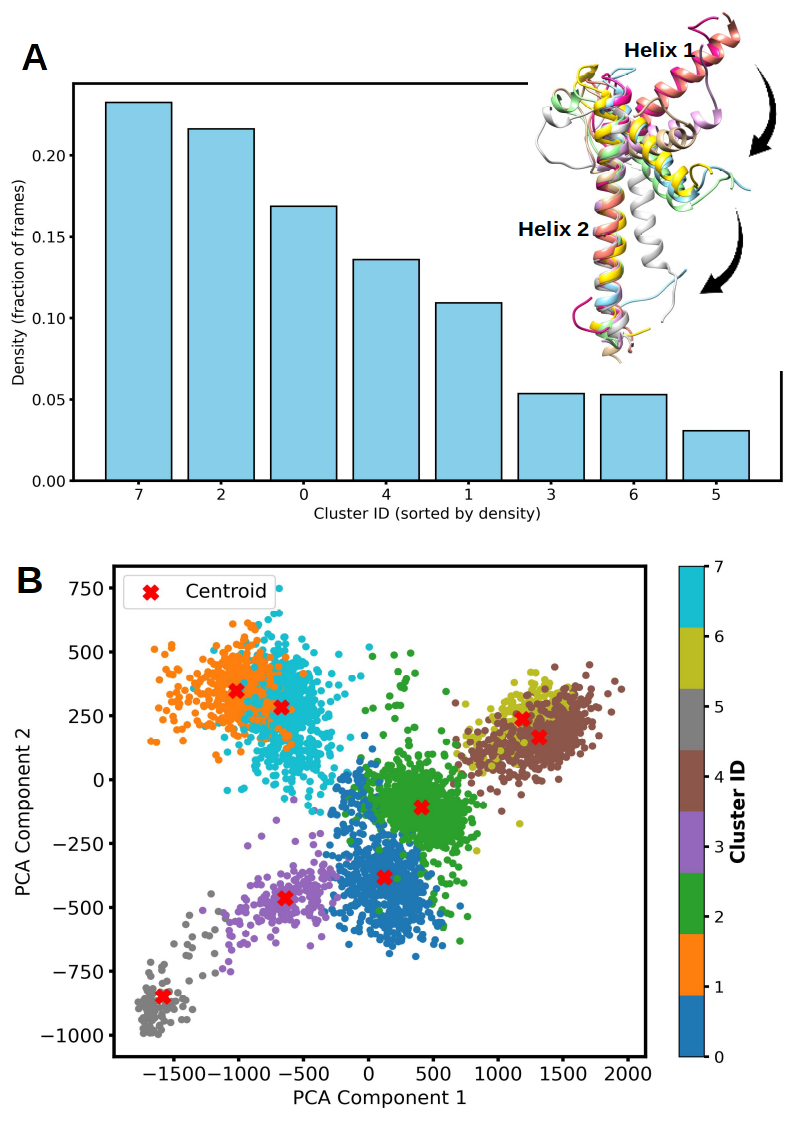

### SI-fig3.png

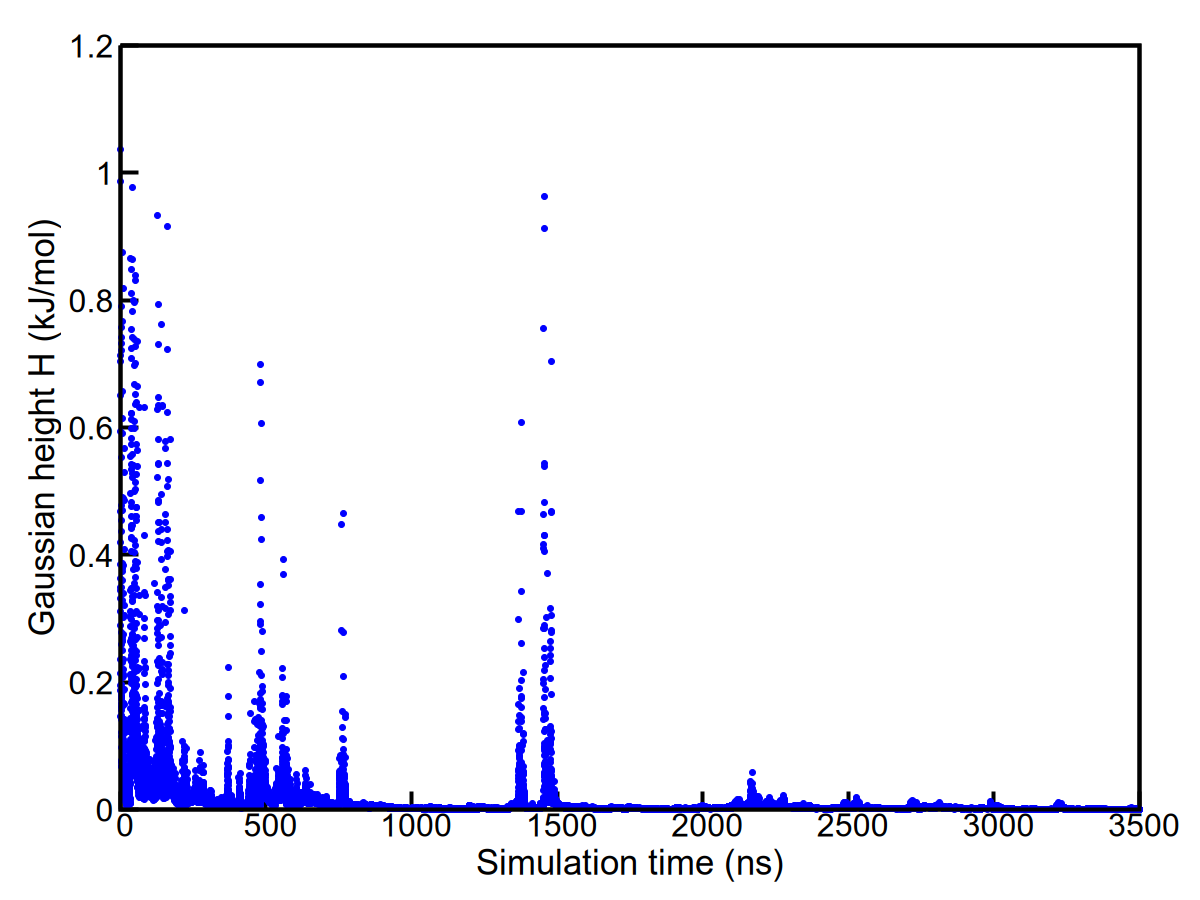

### SI-fig4.png

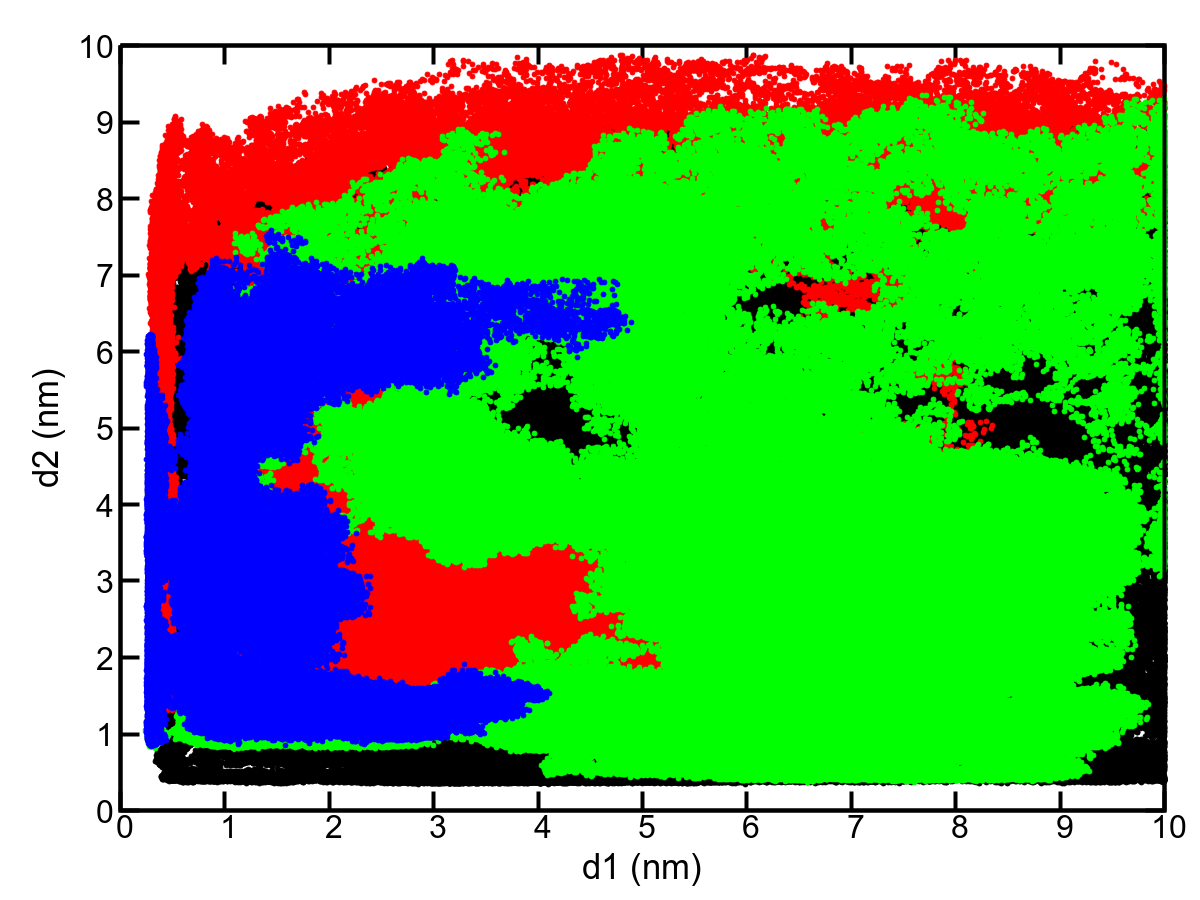

### SI-fig5-a.png

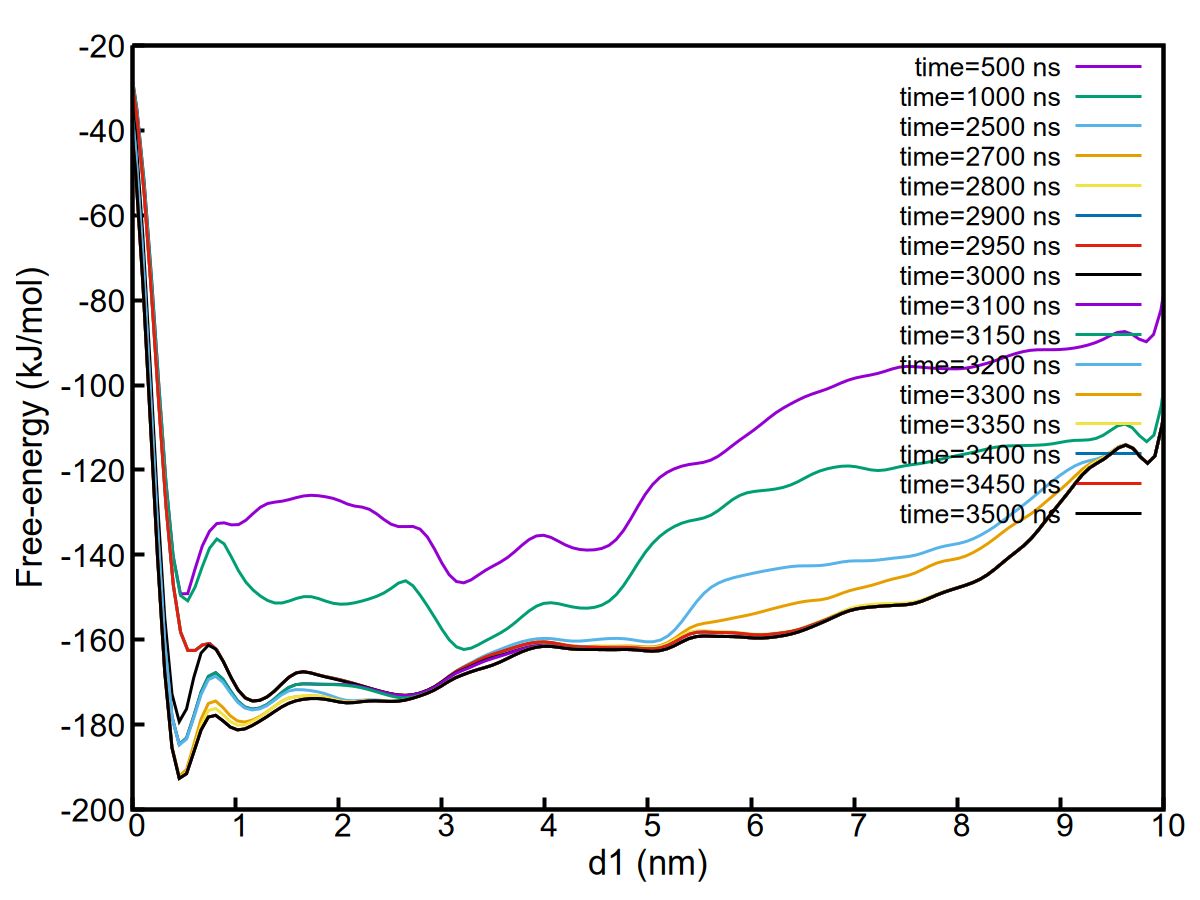

### SI-fig5-b.png

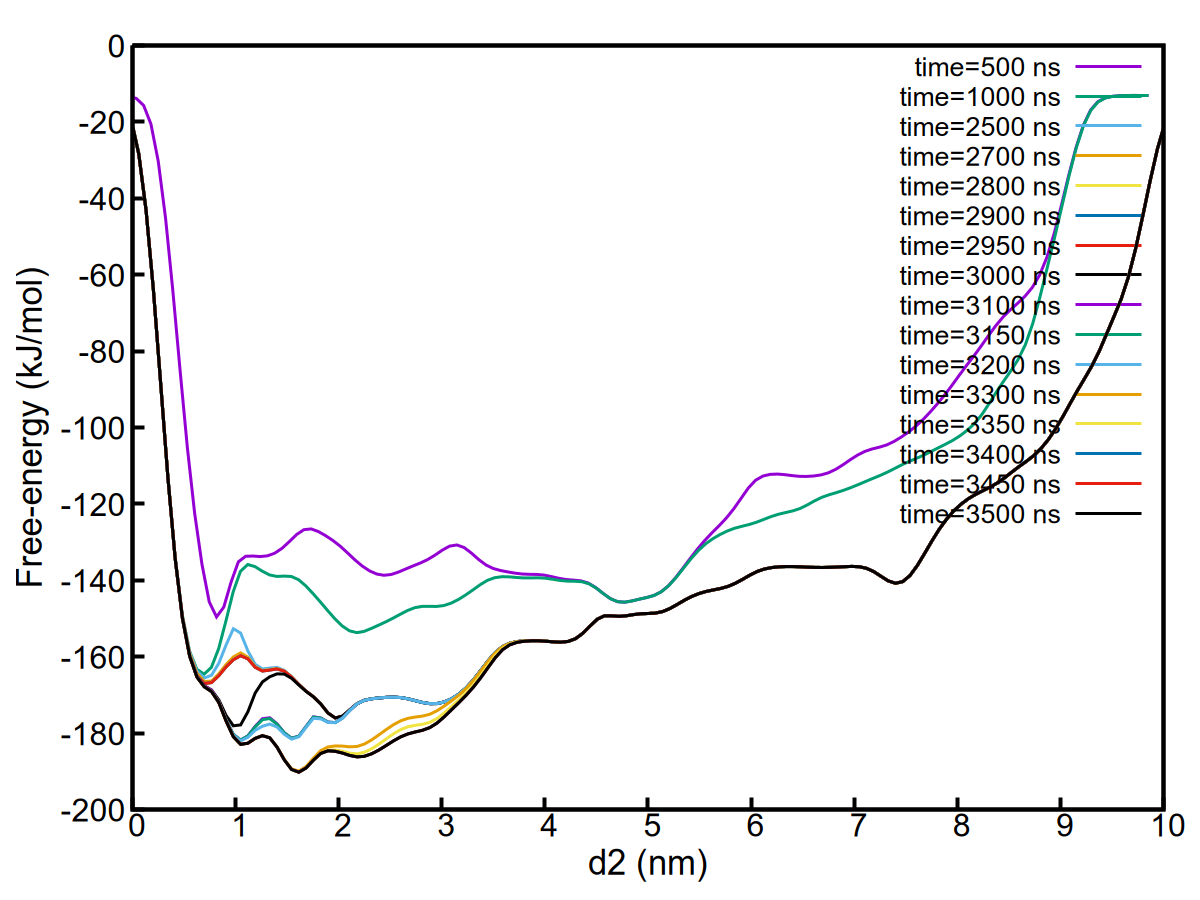

### SI-fig6.png

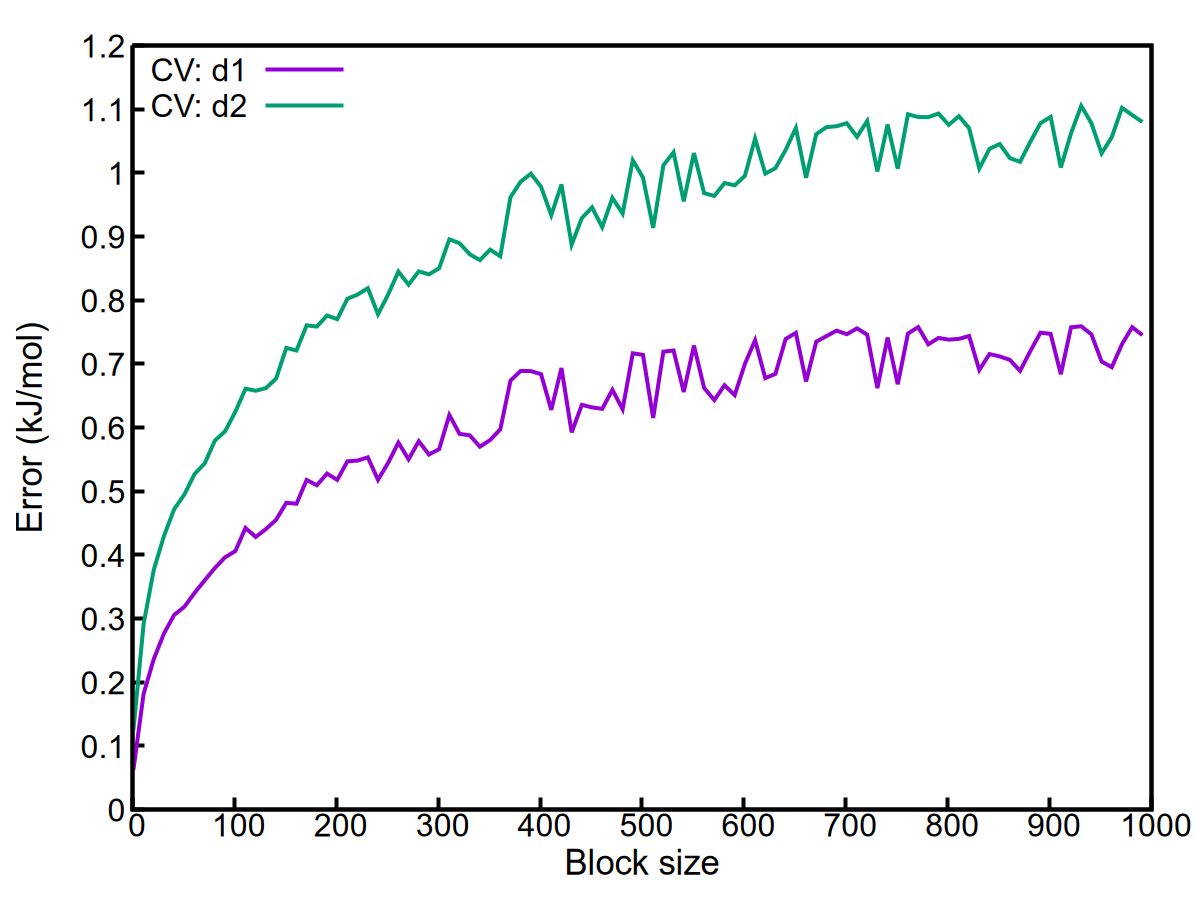

### SI-fig7.png

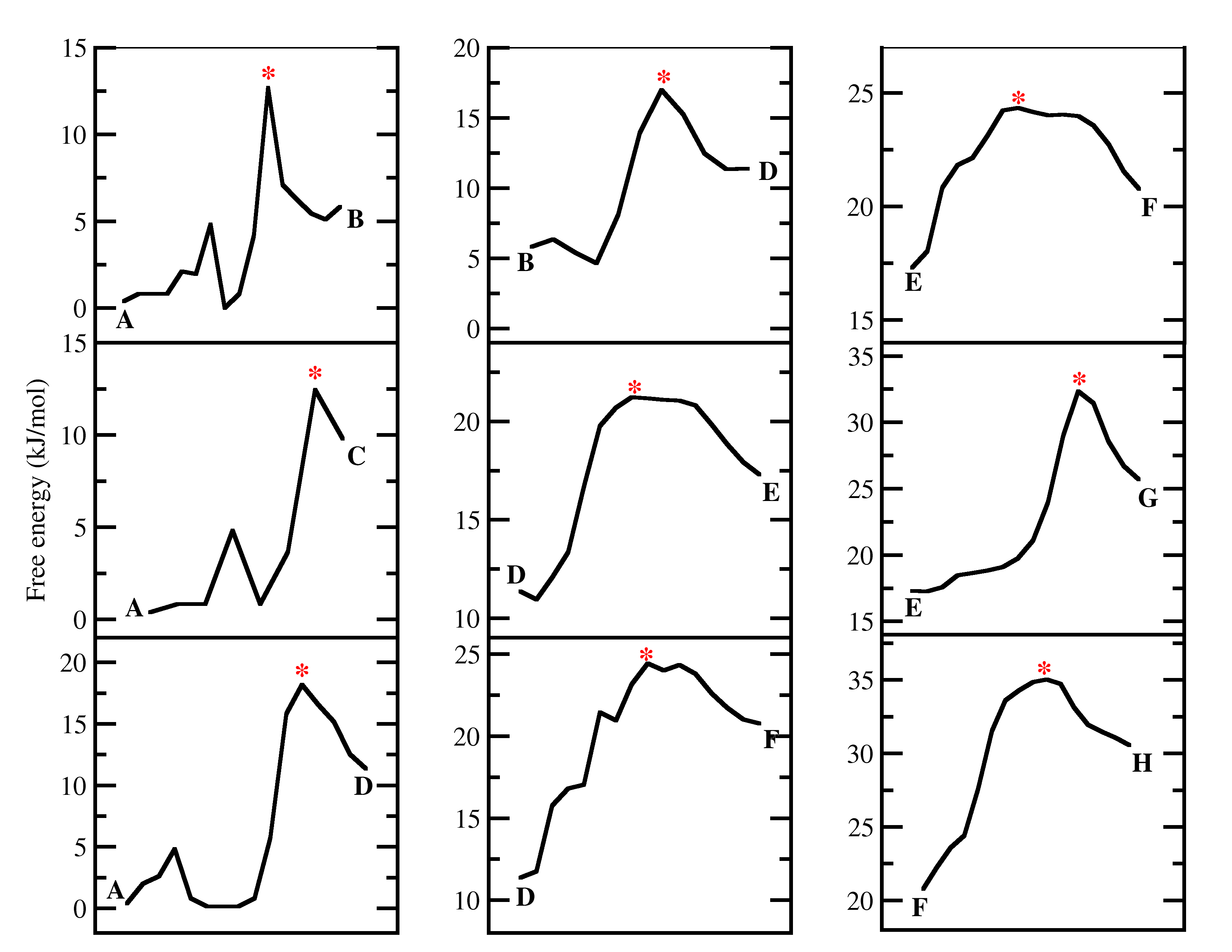

### SI-fig8.png

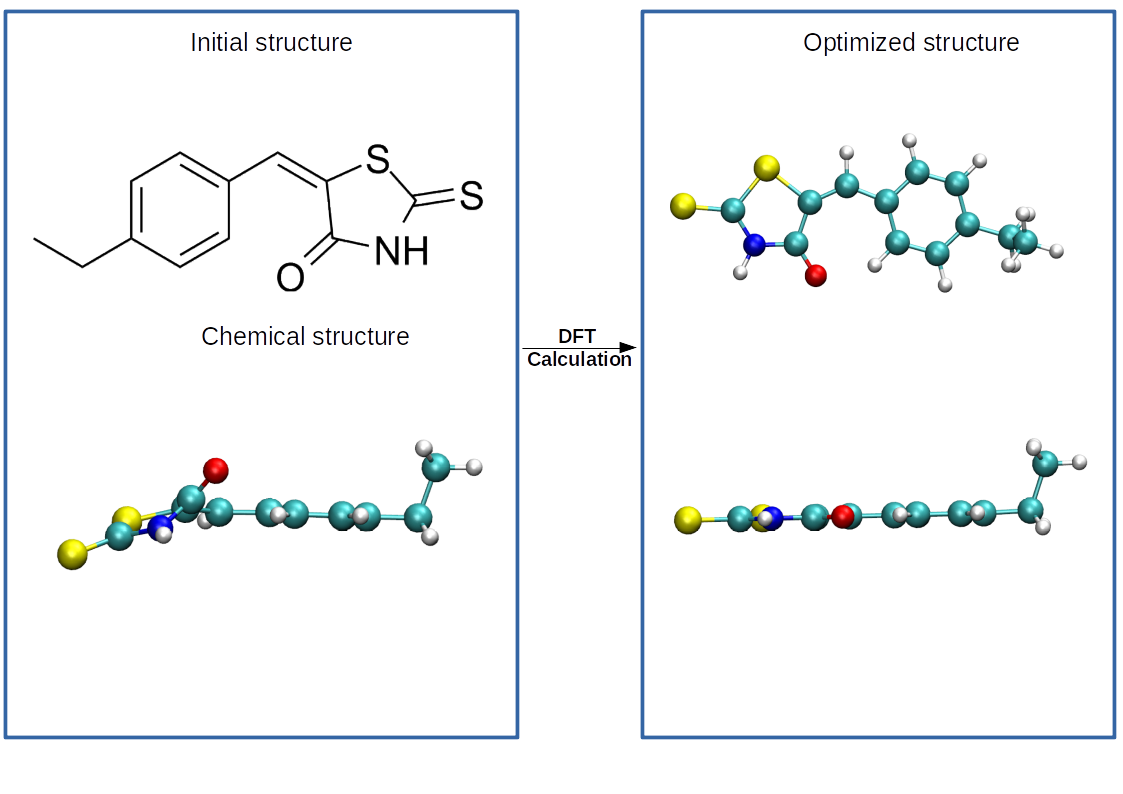

### SI-helicity.png

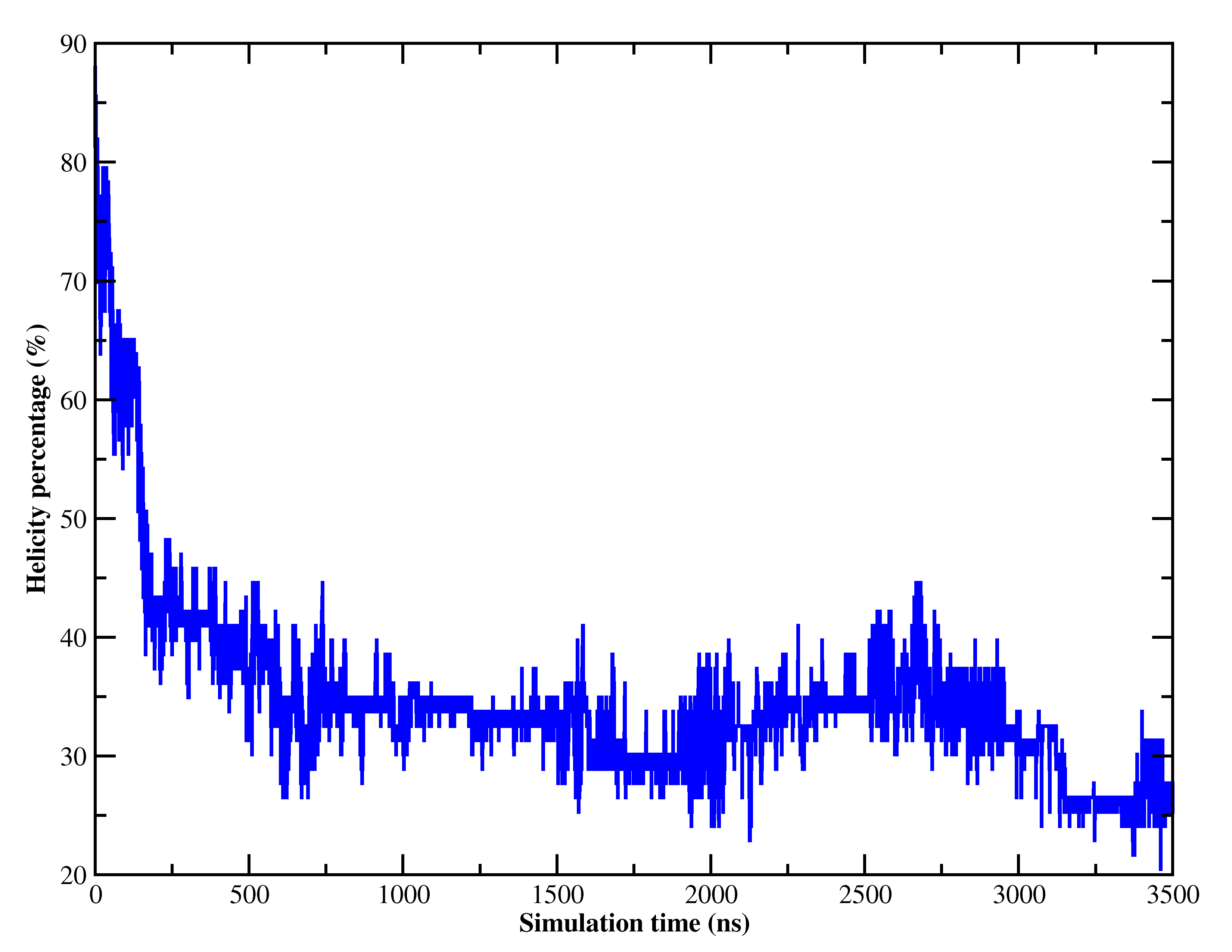
